# HDACs repress runaway stress and cell identity to promote reprogramming in root regeneration

**DOI:** 10.1101/2024.01.09.574680

**Authors:** Ramin Rahni, Bruno Guillotin, Laura R. Lee, Indie Suresh, Brandon M.l. Gorodokin, Graeme Vissers, Kenneth D. Birnbaum

## Abstract

The widespread regenerative capacity of plants is mediated by the ability of specialized cells to reprogram their fate, but the sequential cellular states of regenerating plant cells remain an open question. Here, we characterize the trajectory of cellular reprogramming using single-cell RNA/ATAC-seq, imaging, and mutant analysis. The earliest events were dependent on repressive chromatin modification, where Multiome and genetic analysis showed that Class I histone deacetylases (HDACs) *HDA9* and *HDA19* were needed to shut down old identities and to prevent a runaway stress response. Cell division mediates a second step needed for the acquisition of many new identity markers, where division rates were tuned by DOF transcription factor *OBP1* accelerating and *SMR5*, *7*, and *10* decelerating division rates hours later. The results show how plants actively mediate the loss of remnant identities within hours of injury and then tune cell division rates to rapidly reprogram cells to new identities.

## Introduction

The regenerative capacity of plants has frequently been attributed to their facile ability to dedifferentiate. The concept of dedifferentiation as it was initially conceived necessarily implied active cell division (emphasis ours): “the release of living cells from the confines of their previous organization with *accompanying active mitosis* of these cells” ^1^. Still, even induced callus in Arabidopsis, which was thought to be a dedifferentiated mass of cells, was shown to contain specific cell types ^2^. And studies in axolotl have shown that germ layers retain their identity in the regenerative blastema, another mass of cells thought to be dedifferentiated ^3^. This led Meyerowitz and Sugimoto to point out that the cell states that define the trajectory of dedifferentiation remain largely a black box ^2,4^. In charting the trajectory of regenerating plant cells, chromatin modification and cell division are central in mediating the progression of reprogramming. Thus, defining their roles during the process is central to understanding plant regeneration.

Several lines of evidence show that histone deacetylases (HDACs), which remove acetyl groups on chromatin to increase compaction and repress gene expression, are important in regeneration. In Arabidopsis, treatment with the HDAC inhibitor trichostatin A (TSA) inhibits callus formation from leaf explants, as do mutations in the Class I HDAC *HDA9* or the HD-tuin family member *HDT1* ^5^. Another recent report showed that TSA inhibited shoot regeneration in Arabidopsis at high concentrations but promoted regeneration at low concentrations ^6^. Mutant analysis also showed that *HDA19*, another Arabidopsis Class I HDAC, also promotes *de novo* shoot regeneration from calli ^7^. In rice, plants treated with TSA also show reduced callus formation, as do mutants for the HDAC *OsHDA710* ^8^. By contrast, TSA treatment in wheat has dose-dependent effects, with lower doses promoting callus formation and higher doses having an inhibitory effect ^9^. Chromatin regulation at the histone level has also been implicated in zebrafish fin regeneration, with deacetylation of histone tails via the activity of histone deacetylase 1 ^10^ and demethylation of a repressive H3K27me3 mark ^11^ both being required. Thus, while some inhibition of HDACs may promote organ-specification genes, HDACs appear to be essential for cells to reprogram. HDACs and HATs are additionally known to have cell cycle phase-dependent roles ^12^, although HDACs and HATs also have roles in post-mitotic cells ^13–15^.

Cell division is needed in many different contexts in plants and animals for regeneration to occur^16,17^. In the root epidermis, where hair and non-hair cells form an alternating pattern ^18^, the transcription factor *GLABRA2* (*GL2*) is expressed in differentiating non hair cells ^18^. Interestingly, if hair cells are displaced from a hair file into a non hair file, they change identity, in part by actively reorganizing chromatin around the *GL2* locus during G_1_ ^18^. We recently showed that the cell cycle is accelerated during regeneration and that this acceleration is driven by a shortened G_1_ phase ^19^. A similar dependence of cell fate on division is seen in the floral meristem, where terminal differentiation of floral stem cells relies on eviction of Polycomb group proteins through dilution by division ^20^. Cell division can also allow cells to differentially partition cell fate determinants, such as in the stomatal lineage in leaves ^21^. There, unequal distribution of the protein BASL is associated with early asymmetric divisions. Later, nuclear vs peripheral localization of BASL in the daughters of these divisions correlates with specific cell fates, such as continued asymmetric cell division or terminal differentiation ^21^. Thus, cell division represents an opportunity for a cell to change its fate through chromatin reorganization, partitioning of cytoplasmic determinants, or other processes.

These key processes in regeneration are highly dependent on the plant’s wound response. For example, *WIND1* ^22^, a key regulator of cellular dedifferentiation, and *ERF115* ^23^, a regulator of wound-induced cell division, are both upregulated by wounding. In addition, wounding triggers hormone responses that have been shown to be necessary for regeneration, including auxin ^24^ and jasmonic acid signaling ^25,26^. The mechanical stress of wounding as well as the immediate release of reactive oxygen species are also important triggers for regeneration ^27–29^. However, it appears increasingly likely that wounding also triggers responses that are detrimental to regeneration. For example, we found previously that knocking out glutamate receptors that mediate wound-induced calcium signaling in defense pathways improved regeneration ^30^. Other studies have shown a similar balancing mechanism between defense and regeneration ^31^. Thus, it is not clear whether regeneration requires active mechanisms to suppress an over-reaction to injury–a runaway stress response–to enable regeneration, much like the need to suppress metazoan immune responses to facilitate tissue repair over scarring ^32^.

In addition, we know little about how division and chromatin remodelling mediate the trajectory of a plant cell as it reprograms from one fate to another. How does a cell’s prior fate get erased? For example, the loss of cell identity could be simply mediated by a temporary absence of instructive signals. Alternatively, active mechanisms may be needed to dismantle already specified cell fates. We also do not know if cells must lose their prior fate in order to reprogram to a new cell fate. In the root regeneration system, we know that cells must divide to regenerate ^17^, but we do not know if division is necessary to specify a new cell fate.

Here we map the trajectory of cells in Arabidopsis root tip regeneration—from the loss of remnant identities to the gain of new identities—in a 36-hour span. We show that the loss of cell identity is actively mediated by HDACs, where just a one-hour perturbation of HDACs after injury significantly perturbed root-tip regeneration. In the first two hours, another key role of HDACs was also to rapidly shut down a strong stress response instigated by injury. Mutants in HDACs shown to suppress environmental stresses impaired regeneration, while mutants in plants that promoted the same environmental stress improved regeneration. This sheds light on the concept of a runaway stress response, where at least some of the wound response needs to be suppressed for regeneration to proceed. In addition, cells could embark on reprogramming without division, losing much of their prior identity and expressing early developmental markers. However, when cell division was blocked, cells could not complete reprogramming, failing to express many later, cell-type specific markers for new cell fates. Furthermore, division rates were tightly regulated in a “pump-the-gas, step-on-the-brakes” sequence with *OBF binding protein 1* (*OBP1*) first speeding up cell division rates and members of the *SIAMESE-RELATED* (*SMR*) cyclin dependent kinase inhibitors slowing down division rates in succession. Overall, the reprogramming plant cells first embark on regeneration by actively shutting down prior cell identities and components of the wound response within the first two hours after injury. The cells then express markers for early meristematic “character,” many of which are independent of cell division or histone acetyl group remodelling. In mid stages, rapid divisions allow cells to express mature markers for new cell identities, completing a trajectory from one specialized cell to another within 36 hours.

## Results

### Identification of a regeneration-specific transcriptional program

Arabidopsis roots are able to rapidly replace lost or damaged tissues, even when the entire tip of the root—containing QC, columella, lateral root cap (LRC), and all initials—is surgically removed (**Figure 1a**). Root tip regeneration can be categorized into three stages based on marker analysis: 1) ectopic activation of several stem cell niche markers, 2) loss of remnant identities, and, 3) gain of new identities (**Figure 1b**)^33^. Based on prior studies and our observations, we define a region above the cut stump forming a dome with the height of around 10 cells to be the zone of cellular reprogramming (henceforth the reprogramming zone).

**Figure 1.**
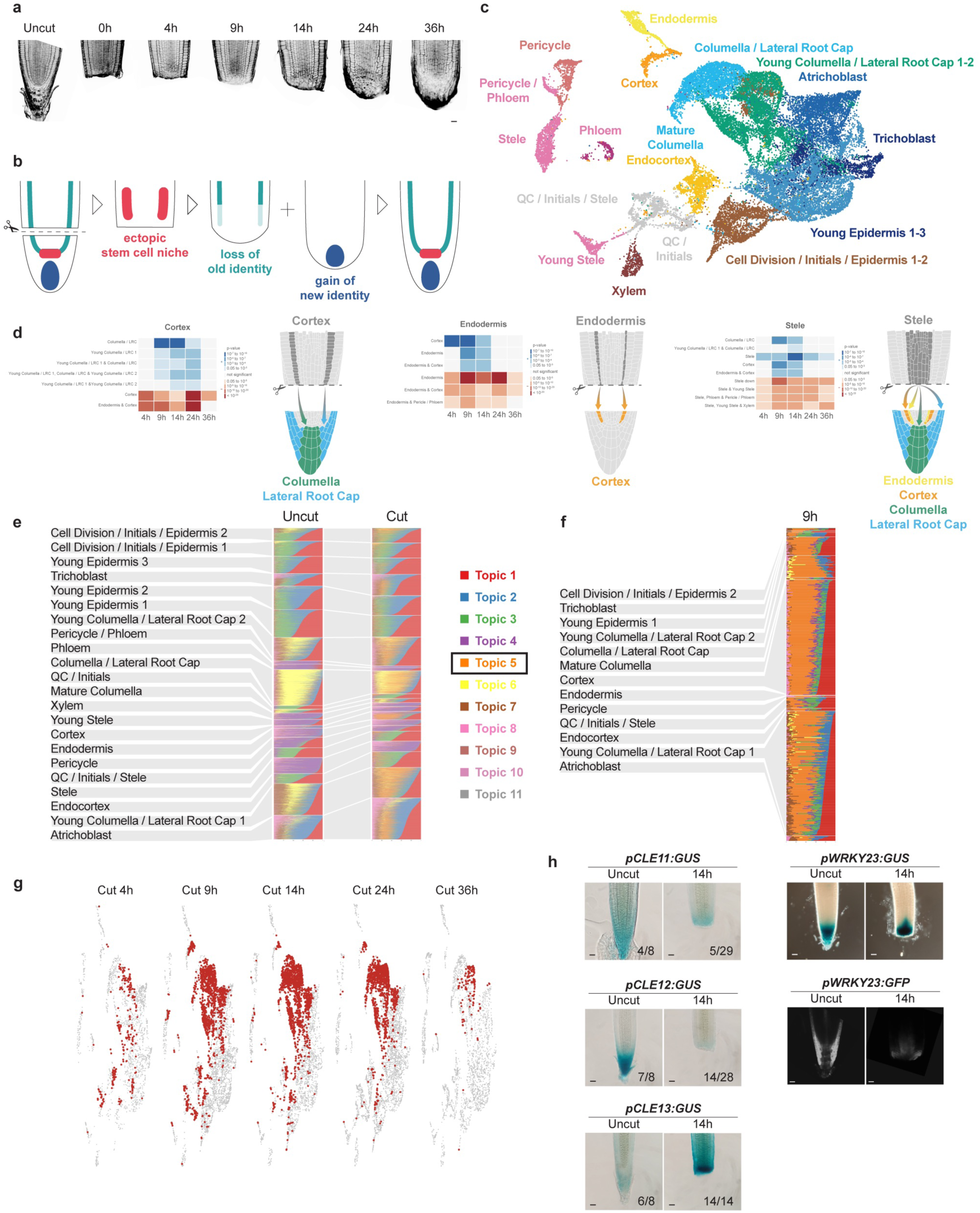
Topic modeling identifies cells undergoing regenerative reprogramming. (a) Confocal time series of a regenerating root with mPS-PI staining. Scale bar corresponds to 20 µm. (b) Schematic showing the three stages of regeneration: ectopic activation of stem cell niche markers (red), loss of old identity (green), and gain of new identity (blue). c) UMAP of uncut root with different cell types in different colors. (d) Heatmaps and schematics showing for cortex, endodermis and stele clusters the differentially expressed genes hypergeometric enrichment with cell type specific markers. In blue upregulated genes and in red downregulated genes compared to 0h. (e) Topic modeling reveals a distinct regeneration-specific identity, in orange (topic 5). (f) A subset of the topic modeling analysis focusing on the 9h timepoint. (g) UMAP showing the distribution over time of cells identified as in regeneration with a high orange topic expression. (h) GUS staining of uncut and 14 hour post cut roots showing the localization of *CLE11, CLE12, CLE13,* and *WRKY23* gene expression, along with confocal images of *pWRKY23:GFP* expression at the same time points. GUS stained roots are unscaled images—scale bar estimated from root length and correspond to 20 µm.

To study these identity transitions more closely, we performed single cell RNA-sequencing (scRNA-seq) in uncut, 4, 9, 14, 24, and 36 hours post cut roots. In total, 30,657 cells were sequenced and combined using the R package Seurat ^34^. Cells from uncut roots (0h) of the same age were used as a reference dataset for integration using SCTranform approach ^34,35^. This approach forces each cell from the cut condition to align onto the uncut reference according to their transcriptomes. The resulting clusters were assigned cell identities based on multiple cell type-specific markers from our own data as well as prior single-cell RNA-seq analyses (**Figure 1c, Supplemental Figure 1a, Supplemental Table 1**).

We used our scRNA-seq regeneration time course data to identify differentially expressed genes between cut and uncut conditions in each cell identity cluster over time in order to map the loss and gain of cell identities. We found that many cell types, such as stele, endodermis or cortex start to lose part of their identities at 9, 14, and 24 hours (**Figure 1d, Supplemental Table 1**). At the same time, these clusters show strong signals of other identities—often belonging to their outer neighbors—at 4 or 9 hours (**Figure 1d**)—consistent with our previously published clonal analysis ^33^. For example, remnant stele cells give rise to adjacent pericycle and endo-cortex, pericycle gives rise to neighboring endodermis, endodermis gives rise to cortex, and the newly formed columella is derived primarily from remnant stele and ground tissue cells (**Figure 1d, Supplemental Table 1**). The signatures of the new identities were lost by 24 to 36 hours, presumably as those cells completed their reprogramming and were indistinguishable from stereotypical cell clusters. Whereas we did not previously find a significant contribution of remnant epidermis to the regenerating root tip ^33^, transcriptional data in the current data suggests that remnant epidermis does in fact contribute to the formation of newly formed columella and lateral root cap at the late 36 hour time point (**Supplemental Table 1**).

To identify a transcriptomic signature unique to regeneration, we used a topic modeling approach ^36^. In this approach, each cell’s transcriptome can be summarized as a combination of K number of topics in varying proportions, where each topic is composed of hundreds of genes that are coexpressed. We performed topic modeling analysis on cells in both cut and uncut conditions using the R package CountClust ^37^ and a total of K=11 topics. Of these 11 topics, one— orange topic (N°5) —was highly enriched in cut cells and almost totally absent from uncut cells (**Figure 1e, Supplemental Figure 1b-d**). In total, we found 3,677 cells with a high proportion of orange topic, compared to only 130 cells in the uncut, representing a false positive rate of 3.4%.

To identify the tissues from which orange topic -enriched cells originated, we looked at their distribution within the integrated UMAP (**Figure 1f,g**). Interestingly, at 4 hours, the orange topic is enriched in clusters annotated as a mix of closely related *QC*, *initials*, *endo-cortex, and epidermis* cells, largely through broad markers of the early meristem, e.g. *PLT1*, *PLT2*, *BBM*, *RGF2*, and *RGF3*. By 9 and 14 hours, this topic is enriched in the *cortex* and *young columella* clusters, the first signs of cellular differentiation. And by 24 and 36 hours, it is more restricted to cells annotated as *columella* and *LRC* (**Supplemental Table 2**), a near complete respecification.

To validate that the orange topic represented genes expressed in the reprogramming zone, we examined five markers that were highly specific to it: *CLAVATA-LIKE*(*CLE*)*11*, *CLE12*, *CLE13*, *WRKY23*, and *OBF binding protein 1* (*OBP1*), with GFP/RFP or GUS reporters. Four showed expression in the reprogramming zone by 9 hours, with *CLE12* showing faint but visible expression (**Figure 1h**). In addition, the orange topic included *PLETHORA1* (*PLT1*), *PLT2*, *SCARECROW* (*SCR*), *BABYBOOM* (*BBM*), *ROOT GROWTH FACTOR 2* (*RGF2*), and *ROOT GROWTH FACTOR 3* (*RGF3*) ^38^ (**Supplemental Table 2**), where *PLT1*, *PLT2*, and *SCR* were previously shown to be induced in the reprogramming zone ^17^. Notably, all these genes are expressed in the early meristem in normal development. However, *OBP1*, and many other markers in the orange topic , were uniquely expressed during regeneration in the reprogramming zone. This showed that the orange topic was composed of both early meristem markers and regeneration-specific genes. Interestingly, the orange topic contained many stress response genes that were not expressed in the uncut root but induced and then rapidly repressed by 14 hours, with this uniquely expressed set of genes making them distinguishable from any cell type in uncut roots. We explore the regulation and the role of these transiently expressed genes below.

Overall, the data show that cells of the proximal meristem left over after the cut rapidly express functional markers for the very early meristem, in line with a switch to a youthful state. At a slightly later time point, cells in the regeneration-specific cluster already possess new cell character, such as endo-cortex, epidermis, and amyloplast-containing columella cells. Thus, we conclude that, in root regeneration, cells dedifferentiate—not to a specifically youthful cell, such as QC per se, but to a regional pro-meristematic character—and then rapidly transdifferentiate to mature cell types.

### Cell division is required for complete reprogramming

We next investigated the different mechanisms influencing the three main stages of regeneration that we identified: 1. ectopic activation of stem cell niche markers, 2. loss of remnant identities, and, 3. gain of new identities. We previously showed that inhibition of the cell cycle inhibits regeneration ^17^, but it was unclear whether some reprogramming could still occur even in the absence of division. Thus, we first sought to determine whether each stage of regeneration we defined was dependent on cell division.

Beginning with live imaging, we examined the requirement of cell division for reprogramming by crossing different transcriptional reporters into a *35S::H2B-mRFP1* background to visualize every nucleus in the root in tandem with cell identity (**Supplemental Figure 2a-d**). Time lapse imaging showed that the QC-specific gene *WOX5* is ectopically activated in cells prior to mitosis (**Supplemental Figure 2a, Supplemental Movie 1**), suggesting division-independence. On the other hand, more mature cap markers like *PET111, IAA10,* and *WEREWOLF (WER)* only appeared after 1 or more observed mitoses, suggesting that division may be required for full reprogramming into mature columella and lateral root cap (**Supplemental Figure 2b-d, Supplemental Movies 2-4**).

To directly test the role of cell division on reprogramming, we treated regenerating roots with the Cyclin-Dependent Kinase inhibitor roscovitine ^39^, and examined its effect on several fluorescent identity markers at 4, 18, and 24 hours post cut. In tandem, we performed scRNA-seq on treated roots at 14 hours post cut—this time encompasses loss of old identity, gain of new identity, and at least one cell division.

First, we tested whether the ectopic reactivation of stem cell niche markers (stage 1) requires cell division. We found, both transcriptionally and using a fluorescent reporter, that *WOX5* expression is still able to return after cell cycle inhibition with roscovitine at 4 (to a limited degree), 14, and 24 hours (**Figure 2a, b**). Another marker of the QC (which additionally marks columella and columella stem cells)—*WIP4*—is also expressed ectopically around 9 hours after root tip removal ^33^. Similar to *WOX5*, *WIP4* expression is detected, albeit at a much lower level, at 24 hours post cut after treatment with roscovitine and was lowly detected in our scRNAseq data 14 hour post cut, suggesting that this marker is more affected by cell cycle inhibition but still comes back at later time points (**Figure 2a, b**). The enhancer trap line *J2341*, marking the initials surrounding the QC, is also typically expressed ectopically between 9-12 hours after root tip removal and returns weakly after cell cycle inhibition (**Figure 2a**). Stem cell niche markers therefore appear to be division-independent. Analysis of scRNA-seq data from roscovitine-treated roots showed that 40% of induced genes were still induced and 70% were still repressed when the cell cycle was blocked, with functional markers of the early meristem, such as *MONOPTEROS* (*MP*) and *PLT2*, showing induction independent of cell division (**Supplemental Table 3**).

**Figure 2.**
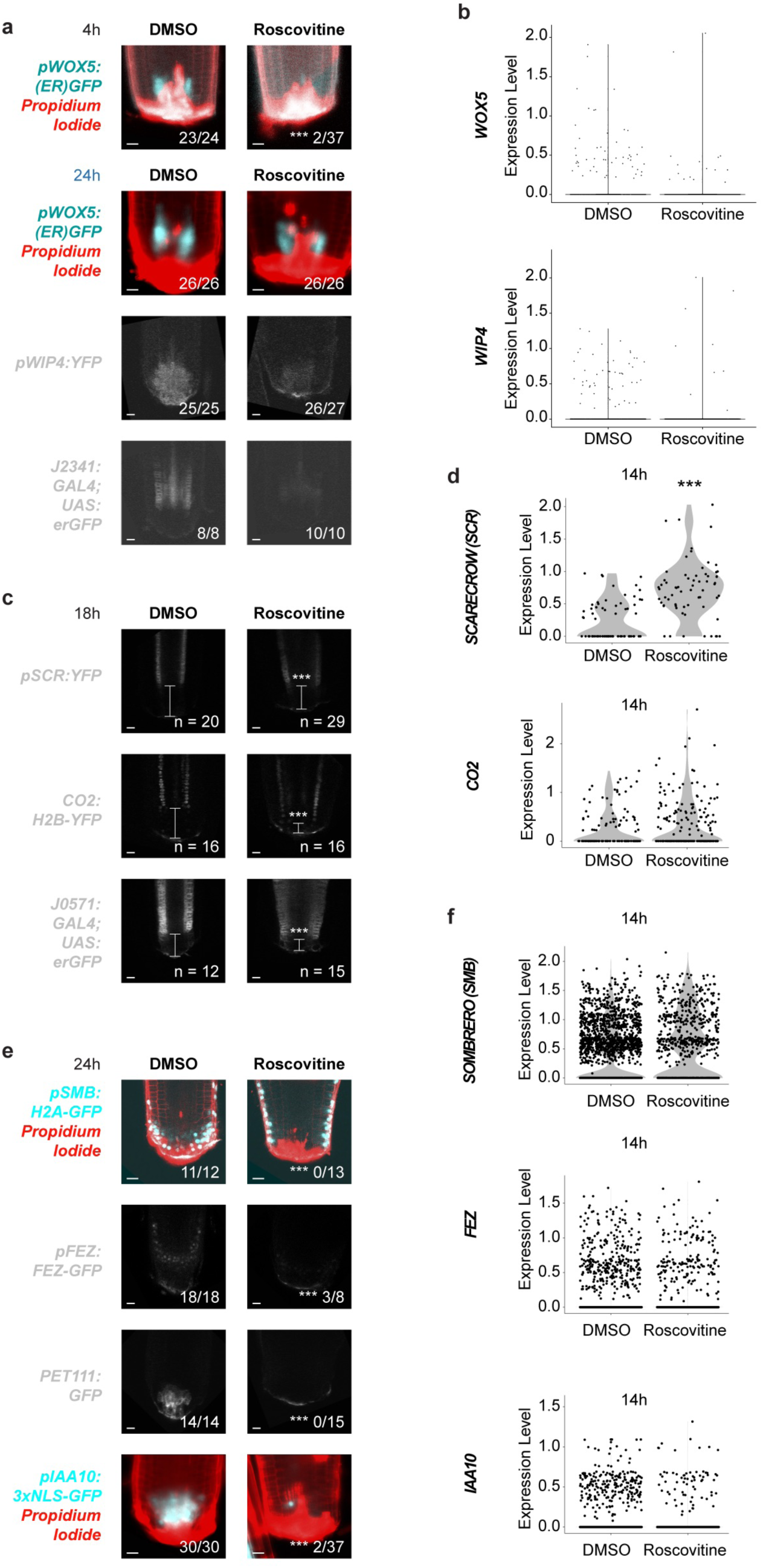
Pharmacological inhibition of cell division perturbs reprogramming. (a) Confocal images of stem cell niche markers *WOX5*, *WIP4,* and *J2341* at 4 and 24 hours post cut with mock (DMSO) or 10 µM Roscovitine treatment. Scale bars correspond to 20 µm. (b) scRNA-seq violin plots showing *WOX5* and *WIP4* expression at 14 hours post cut in mock-(DMSO) or 10 µM Roscovitine-treated roots. (c) Confocal images of remnant identity markers *SCR*, *CO2,* and *J0571* in cut roots at 18 hours post cut after mock (DMSO) or 10 µM roscovitine treatment. Scale bars correspond to 20 µm. (d) scRNA-seq violin plots showing expression levels of *SCR* and *CO2* at 14 hours post cut after mock (DMSO) or 10 µM roscovitine treatment. (e) Confocal images of columella and lateral root cap markers *SMB*, *FEZ, PET111,* and *IAA10* at 24 hours post cut with mock (DMSO) or 10 µM roscovitine treatment. Scale bars correspond to 20 µm. (f) scRNA-seq violin plots showing expression levels of *SMB, FEZ,* and *IAA10* at 14 hours post cut after mock (DMSO) or 10 µM roscovitine treatment. Experiments shown here in Figure 2 were performed at the same time as those in Figure 4 and mock (DMSO) controls for both microscopy and scRNA-seq appear in both. Fractions in a, c, and e indicate frequency of the shown phenotype and asterisks correspond to significance after performing Chi-square with Yates correction. *** = p <0.0001. Scale bars correspond to 20 µm. Asterisks in violin plots correspond to Wilcoxon test between the two conditions *** = p <0.0001.

Next we examined the requirement of cell division on the loss of remnant identities (stage 2). Immediately after root tip removal, markers of remnant tissues (i.e. stele, ground tissue, epidermis) extend to and are flush with the wound. But by 18 hours expression is diminished at the distal-most tip of the root, suggesting cells in this region have lost their original identity (**Figure 2c**, DMSO). To directly test if loss of these remnant identities requires cell division, we examined several transcriptional reporters of remnant identity after cell cycle inhibition with roscovitine. At 18 hours, the domains of the endodermal marker *SCARECROW (SCR)*, the cortex marker *CO2*, and the ground tissue marker *J0571* all typically show diminished expression near the wound (**Figure 2c**). Roscovitine-treated roots still showed loss of expression at the tip, although the region of recession was significantly smaller (**Figure 2c**). In our scRNA-seq data, we found that *SCR* was statistically more expressed in roscovitine treatment compared to mock in endodermal cells. This change was more subtle with CO2 in the cortical cells but followed a similar trend (**Figure 2d**). These results show that a crucial step in reprogramming—loss of former cell identity among specialized cells—is partially dependent on cell division.

To directly test the need for cell division in gain of new identities (stage 3), we examined the effects of cell cycle inhibition on the reappearance of cap-specific markers whose domains are completely removed after wounding. Roscovitine treatment fully inhibited return of the cap markers *PET111*, *IAA10*, *FEZ*, and *SOMBRERO (SMB)* (**Figure 2e**). ScRNA-seq data showed similar trends with fewer cells expressing each of the transcriptional markers after roscovitine treatment than after mock, although these differences were not statistically significant (**Figure 2f**). Analysis of scRNA-seq data from roscovitine-treated roots additionally showed that cap-specific genes–such as *BRAVO*, *PLT1*, *TCP20*, *TMO6*, *WRKY21* and *CDF4–*are inhibited in the presence of roscovitine (**Supplemental Table 3**).

Looking across timepoints, we were able to identify 53% regeneration-specific genes that appear unaffected by cell cycle inhibition and 47% genes that required cell division in order to be activated/repressed. Interestingly, about half of regeneration-specific transcription factors, including *MP* and *OBP1,* are still activated in the presence of roscovitine, suggesting that a portion of the regeneration program is cell cycle-independent (**Supplemental Table 3**). Taken together, these results show that some partial reprogramming—namely the ectopic reactivation of stem cell niche markers—can proceed in a cell cycle-independent manner, but to achieve full reprogramming into mature cell types it is necessary to progress through the cell cycle. In addition, the loss of remnant cell identity that characterizes the early stages of reprogramming appears largely independent of cell division.

### Rapid divisions mediate regeneration and are oppositely controlled by *OBP1* and *SMR10*

In tracking cell division events during live imaging of regenerating roots, one interesting feature we noted was that cells were dividing at a very rapid rate compared to the same region of the root during normal growth: cell cycle durations during regeneration are accelerated nearly 3-fold, from roughly 21 hours ^40^ to roughly 8 hours ^19^ (**Supplemental Figure 3a**). We recently showed that these rapid cell divisions were driven by a shortened G_1_ phase and a concomitant gain of new identity, and driven by a burst of the metabolite glutathione after ablation of the root ^19^.

To further identify the genetic networks involved in regeneration, we took a Mini-Ex single cell gene-network inference approach, using each cell type and the cells in regeneration from every time point (**Supplementary Table 2**). This approach infers transcription factors and downstream targets (i.e. regulons) using both coordinated expression and known transcription factor binding site information. In addition to Mini-Ex, genes specific to or enriched in cells labeled as “in regeneration” were identified using Seurat FindMarker function (**Supplementary Table 2**). These analyses together confirmed several transcription factors already known to have roles in reprogramming—*WRKY23* ^41^, *OBP1* (DOF3.4) ^42^, *TMO7* ^33^—and identified several other genes with no previously known roles in regeneration, such as *MYB15*, and *CLE12* (**Supplemental Figure 1e**).

*OBP1* was previously shown to be induced during regeneration while mutant analysis with an *OPB1* fusion to a repression domain showed defects in root tip regeneration after excision ^42^. We sought to test whether *OBP1*–which was induced as early as 4-9 hours in our scRNA-seq and microscopy data (**Figure 3a, Supplemental Figure 3b**), not disturbed by any of our treatments (**Supplementary Table 3**), and acts upstream of cyclin activity^43^–could have a role in modulating division rates and hence the ability of cells to reprogram. We did not detect an *obp1* (SALK_049540) regeneration phenotype in a traditional gravity recovery assay after cutting, as previously reported ^42^. However, to sensitively test reprogramming efficiency, we developed a quantitative amyloplast assay using modified Pseudo Schiff-PI (mPS-PI) staining against starch-containing amyloplasts as a physical marker for columella cells ^44^. Given that amyloplast-positive cells are removed after wounding, any new amyloplast-positive cells would imply full reprogramming, thus serving as a proxy for reprogramming efficiency in general. Compared to wild type, *obp1* mutants had significantly fewer amyloplast-positive cells at 18 hours **(Figure 3b**). In addition, the *obp1* mutant showed decreased mitotic activity compared to wild type (**Supplemental Figure 3c)**. At 9 hours post injury, when *OBP1* is most strongly expressed in wild type, the mutant also showed significantly fewer accumulated EdU-positive nuclei compared to wild type (**Supplemental Figure 3d**). These results, together with previous findings^42^, suggest that reduction in mitotic ability leads to poorer reprogramming, with *OBP1* being a potential upstream regulator of the rapid divisions observed in regenerating tissues.

**Figure 3.**
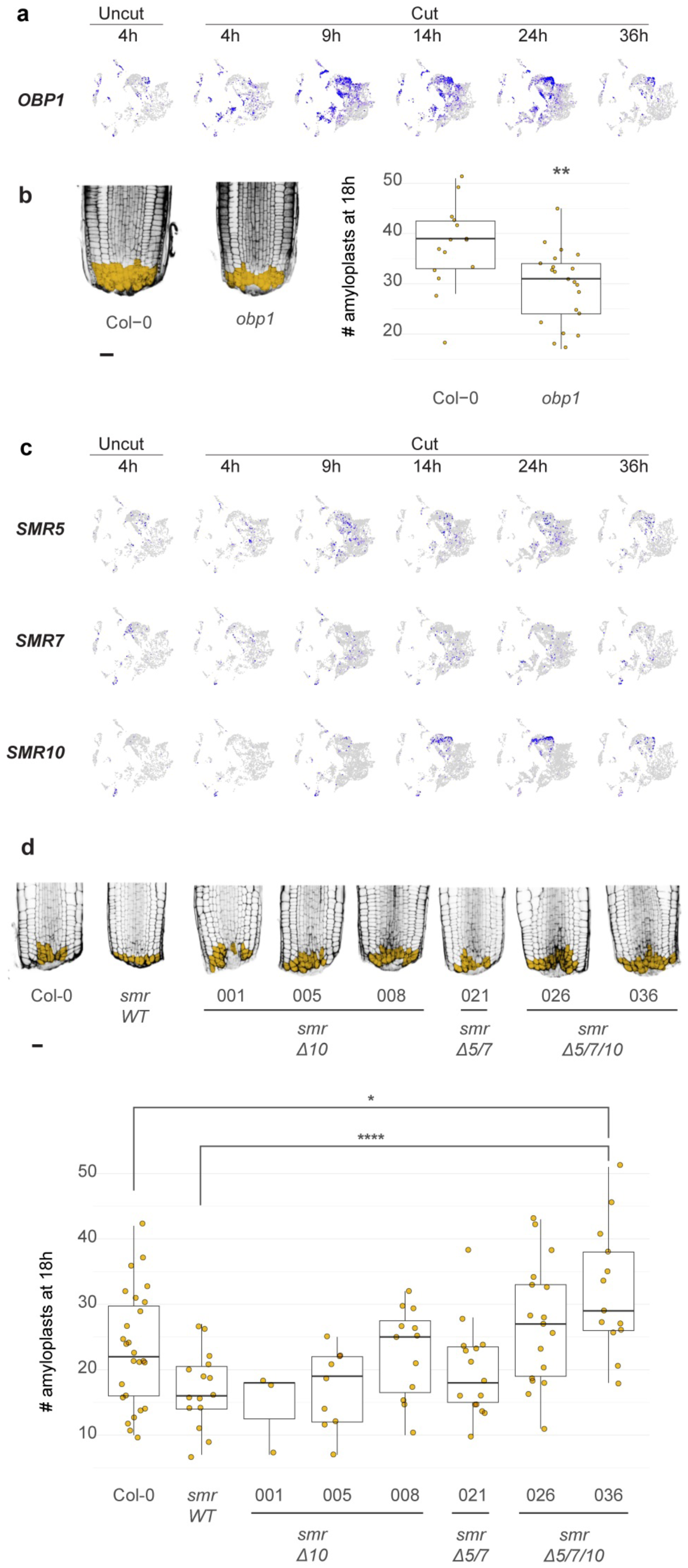
Genetic perturbation of cell cycle regulators affects regeneration efficiency. (a) UMAP of scRNA-seq data in uncut and cut roots at 4, 9, 14, and 24 hours post cut showing broad activation of *OBP1* expression at 9 hours which gradually diminishes at 14 and 24 hours. (b) Boxplot showing number of amyloplast-positive cells at 18 hours post cut in wild type (Col-0) or *obp1* (SALK_049540) mutants as seen by mPS-PI staining. Asterisks indicate significance after t-test: ** = p<0.001. (c) UMAP of scRNA-seq data for *SMR5, SMR7,* and *SMR10* in cut roots at 4, 9, 14, 24, and 36 hours post cut showing activation of *SMR10* at 14 and 24 hours. (d) Boxplot showing number of amyloplast-positive cells at 18 hours post cut in wild type (Col-0) or *SMR* CRISPR mutants. Asterisks indicate significance after t-test: * = p < 0.01, **** = p < 0.00001. Numbers on X axis represent the number of amyloplasts at 18 hours post cut.

By contrast, Cyclin-Dependent Kinase Inhibitors (CDKIs) are negative cell cycle regulators ^45^, and one CDKI, *SIAMESE-RELATED 10 (SMR10),* was highly upregulated in our scRNA-seq dataset in later timepoints, with its close homologs *SMR5* and *SMR7* also showing induction albeit at a much weaker level (**Figure 3c**). We hypothesized that *SMR5, 7,* and *10* may be restoring homeostasis at later time points by suppressing the rapid cell cycles activated during the early stages of regeneration. If so, we predicted that mutants for these CDKIs would have an improved reprogramming efficiency. Using CRISPR-Cas9, we generated an *SMR10* single mutant (*smrΔ10*), a double mutant of its close homologs *SMR5* and *SMR7* (*smrΔ5/7*), and a triple mutant (*smrΔ5/7/10*). Compared to wild type, higher order mutants showed increasing numbers of amyloplast-positive cells at 18 hours post cut, with one *smrΔ5/7/10* allele having significantly more amyloplasts compared to wild type (**Fig 3d**) and increased mitotic activity (**Supplemental Figure 3e,f**). These data suggest that an increase in mitotic ability via inhibition of CDKIs facilitates reprogramming.

The opposite regeneration phenotypes of *obp1* mutants compared to *smr* mutants are consistent with a mechanism wherein *OBP1* is activated relatively early in regeneration to accelerate the cell cycle and facilitate reprogramming, whereas *SMR5*, *7*, and *10* act later to “put on the brakes” and transition cells from a more proliferative state to homeostatic division rates. This shows how the plant finely tunes division rates during regeneration to influence the speed of cellular reprogramming, in keeping with our findings that complete reprogramming of cell identities is dependent on the progression of the cell cycle.

### HDAC and HAT activities are required for complete reprogramming

One mechanism for cell cycle-dependent reprogramming in plants is the modulation of chromatin accessibility, which is regulated—among other mechanisms—by the countervailing activity of histone deacetylases (HDACs) and histone acetyltransferases (HATs). In our topic modeling approach above, we found that the “cells in regeneration” identified by the orange topic , especially those annotated as *QC* / *initials,* were enriched for genes that showed dynamic regulation of chromatin accessibility during regeneration in previous published studies, showing greater accessibility after injury (**Supplemental Figure 4a**) ^46,30^. This led us to speculate that early steps in the reprogramming process could be mediated via regulation at the chromatin level, such as (de)acetylation of histone tails.

To examine the effects of histone acetylation state on reprogramming efficiency, we began with a pharmacological approach using either trichostatin A (TSA), a pan-HDAC inhibitor ^47^ that effects hyperacetylation of histone tails ^48^, curcumin, a specifically inhibitor of the CBP/p300 HAT family ^49^, or MB-3, an inhibitor of the GCN5/GNAT/MYST family of HATs ^50^. MB-3 treatment did not lead to visible effects on regeneration, even at higher concentrations (**Supplemental Figure 4b**). We therefore focused on curcumin treatment for HAT perturbations, which led to partial inhibition of regeneration, and TSA treatment for HDAC perturbations, which completely blocked regeneration. We performed microscopy and scRNA-seq on regenerating roots treated with TSA or curcumin, focusing on the three phases of regeneration outlined in Figure 1b. In addition, we repeated scRNA-seq experiments, cutting roots and transferring them to either TSA or curcumin for 14 hours (when new cell identity markers are induced) and then harvesting cells.

Beginning with the ectopic reactivation of stem cell niche markers, we found that expression of *WOX5* was unperturbed in both TSA- and curcumin-treated roots, showing ectopic activation at both 4 (albeit weakly in TSA) and 24 hours (**Figure 4a**). In the scRNA-seq of TSA- or cucumin-treated regenerating roots, the *WOX5* gene showed a similar induction after cutting, with somewhat lower expression in the curcumin treatment (**Figure 4b**). The extent and localization of the *WOX5* fluorescent domain in TSA-treated roots at 24 hours resembled mock-treated roots around 4 hours, suggesting that HDAC inhibition leads to “stalling” at early stages. By contrast, the *WOX5* domain after curcumin treatment morphologically resembles slightly later mock-treated regeneration stages, with complete regeneration still delayed overall in this treatment. Next we examined *WIP4* expression after treatment with TSA or curcumin and found that, while the marker still returned at a lower expression level after curcumin treatment, its was completely blocked by TSA at 24 hours (**Figure 4a**) and dramatically reduced by both treatments in the scRNA-seq data (**Figure 4b**). This is in contrast to roscovitine, which had a much less dramatic effect on *WIP4* expression (Figure 2a,b), suggesting that while *WIP4* activation is division-independent, it is HDAC-*dependent*. This also importantly shows that roscovitine and TSA have effects that can be uncoupled, and suggests some level of independence between cell division and chromatin remodeling of early regeneration markers.

**Figure 4.**
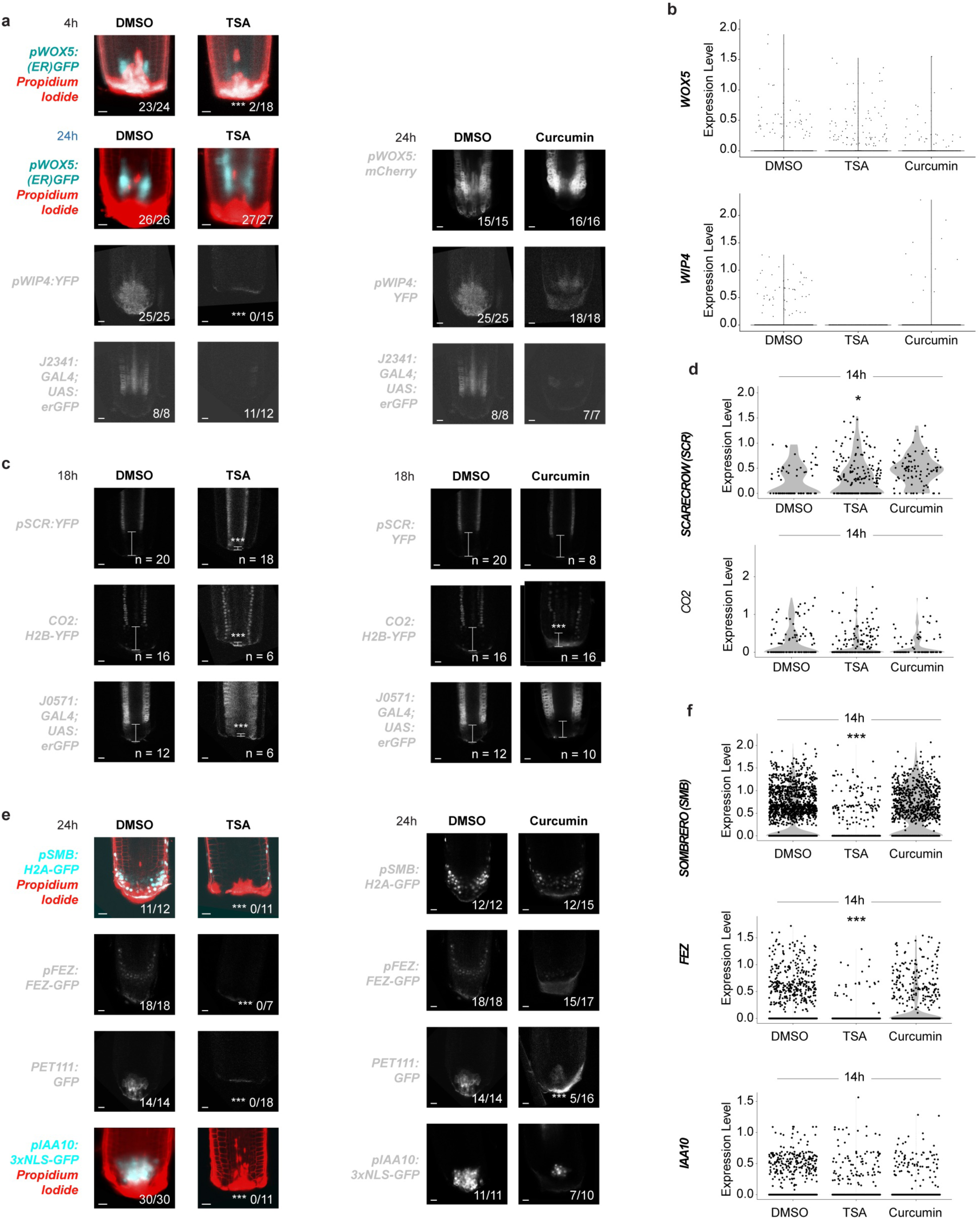
Pharmacological inhibition of HDAC and HAT activity perturbs regeneration. (a) Confocal images of stem cell niche markers *WOX5*, *WIP4,* and *J2341* at 4 and 24 hours post cut with mock (DMSO), 10 µM TSA, or 10 µM curcumin treatment. (b) scRNA-seq violin plots showing *WOX5* and *WIP4* expression at 14 hours post cut in mock-(DMSO), 10 µM TSA-, or 10 µM Curcumin-treated roots. (c) Confocal images of remnant identity markers *SCR*, *CO2,* and *J0571* in cut roots at 18 hours post cut after mock (DMSO), 10 µM TSA, or 10 µM Curcumin treatment. (d) scRNA-seq violin plots showing expression levels of *SCR* and *CO2* at 14 hours post cut in mock-(DMSO), 10 µM TSA-, or 10 µM curcumin-treated roots. (e) Confocal images of columella and lateral root cap markers *SMB*, *FEZ, PET111,* and *IAA10* at 24 hours post cut with mock (DMSO), 10 µM TSA, or 10 µM curcumin treatment. (f) scRNA-seq violin plots showing expression levels of *SMB, FEZ,* and *IAA10* at 14 hours post cut in mock-(DMSO), 10 µM TSA-, or 10 µM curcumin-treated roots. Experiments shown here in Figure 4 were performed at the same time as those in Figure 2 and mock (DMSO) controls for both microscopy and scRNA-seq appear in both. Fractions in a, c, and e indicate frequency of the shown phenotype and asterisks correspond to significance after performing Chi-square with Yates correction. *** = p <0.0001. Asterisks in violin plots correspond to the Wilcoxon test between the conditions, * = p <0.05, *** = p <0.0001. All scale bars correspond to 20 µm.

To examine the extent of the dependence of histone acetylation changes on cell division, we calculated the overlap between all three treatments in scRNA-seq data. Considering genes induced in regeneration in mock treatments, each of the treatments individually impaired the induction of a large percent: 61% (roscovitine), 38% (TSA), 38%(curcumin). The treatments also impaired the repression of genes normally downregulated in mock treatments: 31% (roscovitine), 45% (TSA) and 67% (curcumin) (**Supplemental Table 3, Supplemental Figure 4d)**. However, the results showed moderately high independence in the effects of chromatin remodeling and cell division on global expression. For example, if histone deacetylation was dependent on cell division, we would expect a high degree of overlap between TSA-treated regenerating roots and the roscovitine-treated regenerating roots. However, 330 genes (50%) blocked by TSA treatment were still induced in roscovitine treatments. Similarly, 493 genes (59%) blocked by curcumin treatment were still induced in roscovitine treatments. And 295 genes (44%) blocked by TSA treatment were still induced in curcumin treatments (**Supplemental Table 3).** Overall, this shows that chromatin-level changes during reprogramming, especially acetylation levels, can work independently of division.

TSA has the potential for mildly cytotoxic effects when used *in planta* ^48^. To rule out that TSA-induced failure to regenerate is due to unintended cell cycle inhibition, we combined TSA treatment with EdU labeling. TSA- and mock-treated roots both showed uniform S-phase entry after 24 hours (**Supplemental Figure 4c**). Moreover, cells clearly mid-mitosis were observed in both conditions, suggesting that neither S phase entry nor mitosis are significantly impaired by TSA (**Supplemental Figure 4c**).

To ask whether remnant identity loss was dependent on acetylation state, we treated regenerating roots with TSA or curcumin and examined remnant identity markers *SCR, CO2,* and *J0571,* which typically show dramatically diminished expression near the wound in regeneration (**Figure 4c**, DMSO). Interestingly, all the examined remnant identity markers showed virtually no change in expression near the wound in TSA-treated roots compared to mock, suggesting that these remnant identity genes fail to shut down, in line with the hyperacetylation of histone tails associated with TSA treatment (**Figure 4c**). HAT inhibition via curcumin treatment, by comparison, had a less dramatic effect on loss of remnant identities, with only the cortex marker *CO2* being appreciably diminished near the wound (**Figure 4c**). Similarly, in scRNA-seq analysis, TSA-treated plants retained a significantly greater number of *SCR*-expressing cells compared to mock-treated plants while curcumin had no effect (**Figure 4d**). However, this effect was not statistically different for *CO2* (**Figure 4d**).

Examining the gain of new identities (stage 3), we focused on the return of columella and lateral root cap markers after treatment with TSA or curcumin. In TSA-treated roots, regeneration fails, whereas curcumin-treated roots are able to partially regenerate. To test if these outcomes are due to a failure to specify new cap identities, we examined several distal identity markers in roots treated with either TSA or curcumin. Fluorescent reporters for *PET111, SMB, FEZ,* and *IAA10* all failed to return in TSA-treated roots compared to mock (**Figure 4e**), similar to their failure to be induced in the cell-cycle blocks (Figure 2). Similarly, scRNA-seq analysis of *SMB, FEZ,* and *IAA10* showed that *SMB* and *FEZ* were most significantly inhibited after TSA treatment (**Figure 4f**). Meanwhile, curcumin-treated roots had a reduction in the number of cells gaining these new identity markers during regeneration, in line with partial regeneration (**Figure 4e**).

TSA overall had more dramatic effects on regeneration and affected earlier stages of regeneration—roots appeared “stalled” at an early post-cut morphology, and both loss of remnant identities and gain of new identities was significantly perturbed. By contrast, curcumin primarily affected gain of new identities, a late stage process in regeneration. The effect of TSA on early stage markers that was partially independent of cell division–the rate of which increased by 6-8 hours–was intriguing for the potential of sequential roles of chromatin remodeling and cell division.

### A 1 hour inhibition of HDACs immediately after injury is sufficient to severely impair both loss of cell identity and gain of new identities

Given that we observed early loss of remnant identities at both scRNA-seq and microscopic levels, and that inhibition of HDAC activity blocked both processes, we asked what time point in regeneration was the most crucial for HDAC activity relative to HATs and cell cycle regulators. To do so, we performed a series of pulse-chase experiments on roots by treating them with progressively shorter windows of HDAC, HAT, or cell cycle inhibition, each of which was followed by a chase period on standard growth medium to recover (**Figure 5a**). Regeneration was scored as either “full”, resembling mock-treated roots at 48 hours post cut, “partial”, wherein some amyloplasts are visible via lugol staining but the conical morphology of the root tip is not re-established, “stalled”, which resemble 0 hour post cut roots, and “failed”, where epidermal cells at the tip of the root expand, terminally differentiate, and grow root hairs (**Supplemental Figure 5a**).

**Figure 5.**
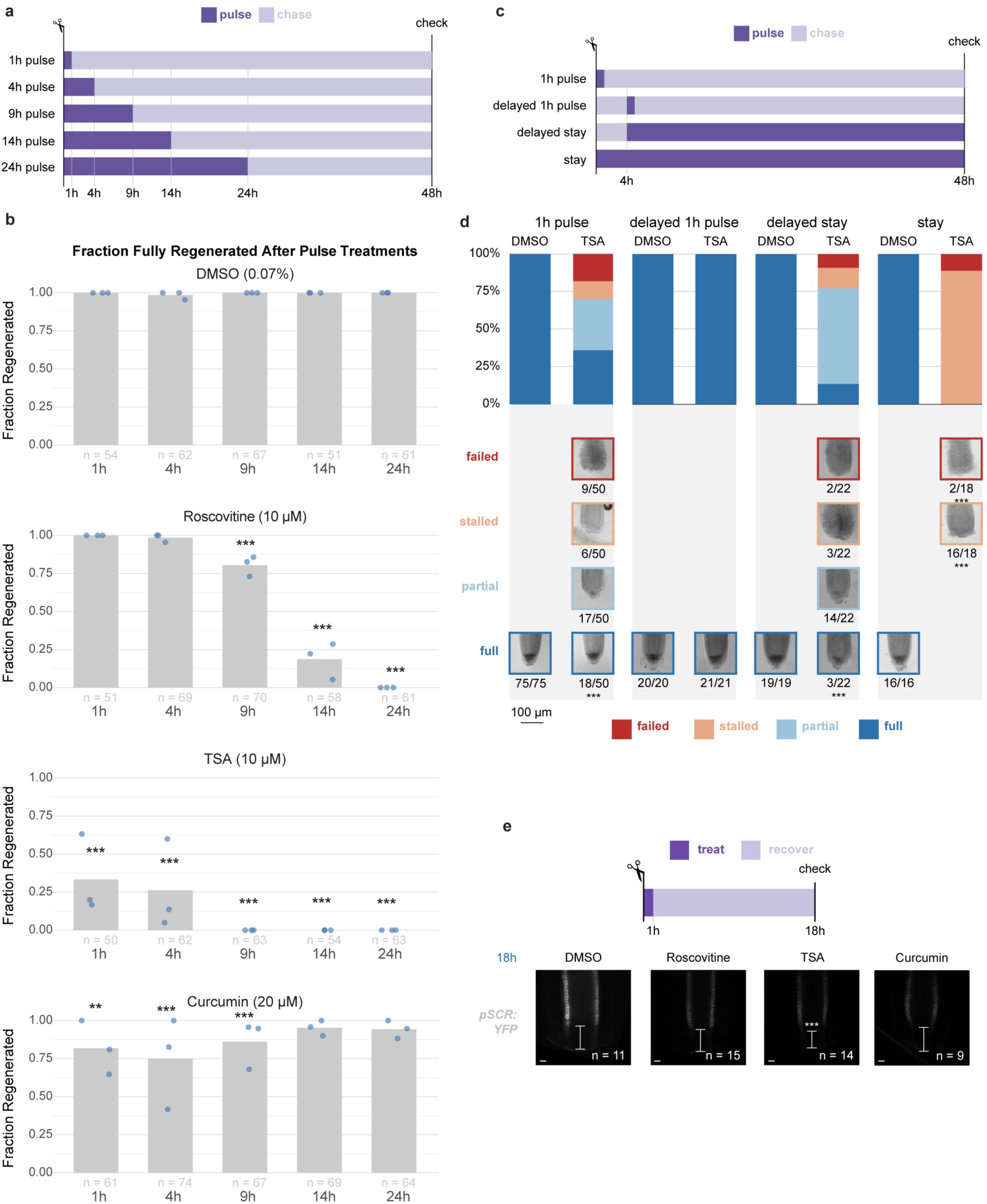
Histone deacetylase activity is needed in the first hour after wounding. (a) Experimental scheme—plants were cut and treated with mock (DMSO), roscovitine, TSA, or curcumin for 1, 4, 9, 14, or 24 hours, before recovery up to 48 hours post cut. (b) Results from the experiments outlined in a: Roscovitine-treated roots start to affect regeneration at 9 or more hours of treatment, whereas TSA treatment has an effect after only a 1 hour pulse. (c) Experimental scheme—plants were cut and given a 1 hour pulse of mock (DMSO) or TSA either immediately after cutting, or after waiting 4 hours post cut. Alternatively, plants were transferred to treatments 4 hours post cut and kept there, or simply kept on treatment media for the full 48 hours as a positive control. (d) Results from the experiments outlined in c: A 1 hour pulse of TSA immediately after cutting significantly reduces the number of fully regenerated root phenotypes, as does a delayed stay or a full stay on the inhibitor. Unscaled images—scale bars estimated from root width and correspond to 20 µm. (e) A 1h pulse of TSA immediately after cutting also affects loss of remnant identities, as seen by less disappearance of *pSCR:YFP* near the cut site compared to mock (DMSO), roscovitine, or curcumin. Scale bars correspond to 20 µm. * = p < 0.01, ** = p < 0.001, *** = p <0.0001

With a 24-hour pulse, TSA and roscovotine almost completely blocked regeneration, while roots treated with curcurmin regenerated almost normally (**Figure 5b**). With a 14-hour pulse, both roscovotine and TSA had severe reductions in the number of fully regenerated roots, but roscovotine-treated roots began to show improvements in regeneration, which recovered to nearly mock levels in the 9-hour pulse (**Figure 5b**). TSA significantly perturbed full regeneration at 9, and 4 hour pulses, and even after just a 1-hour pulse. To rule out that lingering TSA was causing the effects seen by the 1-hour pulse, we cut roots, waited 4 hours, then applied a 1-hour pulse of TSA (**Figure 5c**). This delayed 1-hour pulse had no effect on regeneration, whereas keeping roots on TSA starting 4 hours post cut had a similar effect to the one hour pulse (**Figure 5d**). As above, full treatment for the entire duration of regeneration completely blocked regeneration (**Figure 5d**).

To corroborate that the brief HDAC block could be long enough to affect loss of cell identity, we treated roots carrying the *SCR* marker, which recedes rapidly after injury, with a one hour TSA treatment. Indeed, the one-hour TSA treatment also impaired SCR recession from the tip (i.e., loss of identity) (**Figure 5e**).

The result pointed to a strong role for histone deacetylation in the first hour after injury, followed by strong effects from cell cycle blocks between 9 and 14 hours, while we detected only weak effects for curcumin at all time windows.

### The suppression of stress responses and remnant cell identities are dependent on HDACs

The effects of histone deacetylation inhibition for only one hour were surprisingly early so we sought to examine the consequences of a brief TSA treatment at both the chromatin and mRNA levels by performing single nucleus 10x Multiome sequencing (10x Genomics). The Multiome protocol allowed us to obtain single nucleus RNA-seq (snRNA-seq) and snATAC-seq profiles within the same cell for each condition. Both uncut and cut roots were treated with mock (DMSO) or TSA for 2 hours before processing. We selected a 2-hour treatment window over the previous 1 hour one to give us enough time to reliably process the samples in parallel for better reproducibility. This created four datasets that were integrated together using the R package Signac (see Methods, **Figure 6a, Supplemental Figure 6a-e**), and cell identity was determined using canonical cell type markers on the gene expression dataset (**Supplemental Figure 6f**). Interestingly, we found a cluster—N°6—that was highly specific to the cut datasets. This cluster did not display strong canonical cell type marker expression (**Figure 6a, Supplemental Table 4**), but was enriched for genes related to glutathione response–in line with recently published work showing its role in = rapid cell divisions=^19^. This suggests that cluster N°6 represents the cells immediately responding to wounding. Interestingly, this cluster was present in both of the cut mock and TSA conditions, suggesting that TSA does not fully impair these early changes happening during the first 2 hours after wounding (**Figure 6a**).

**Figure 6.**
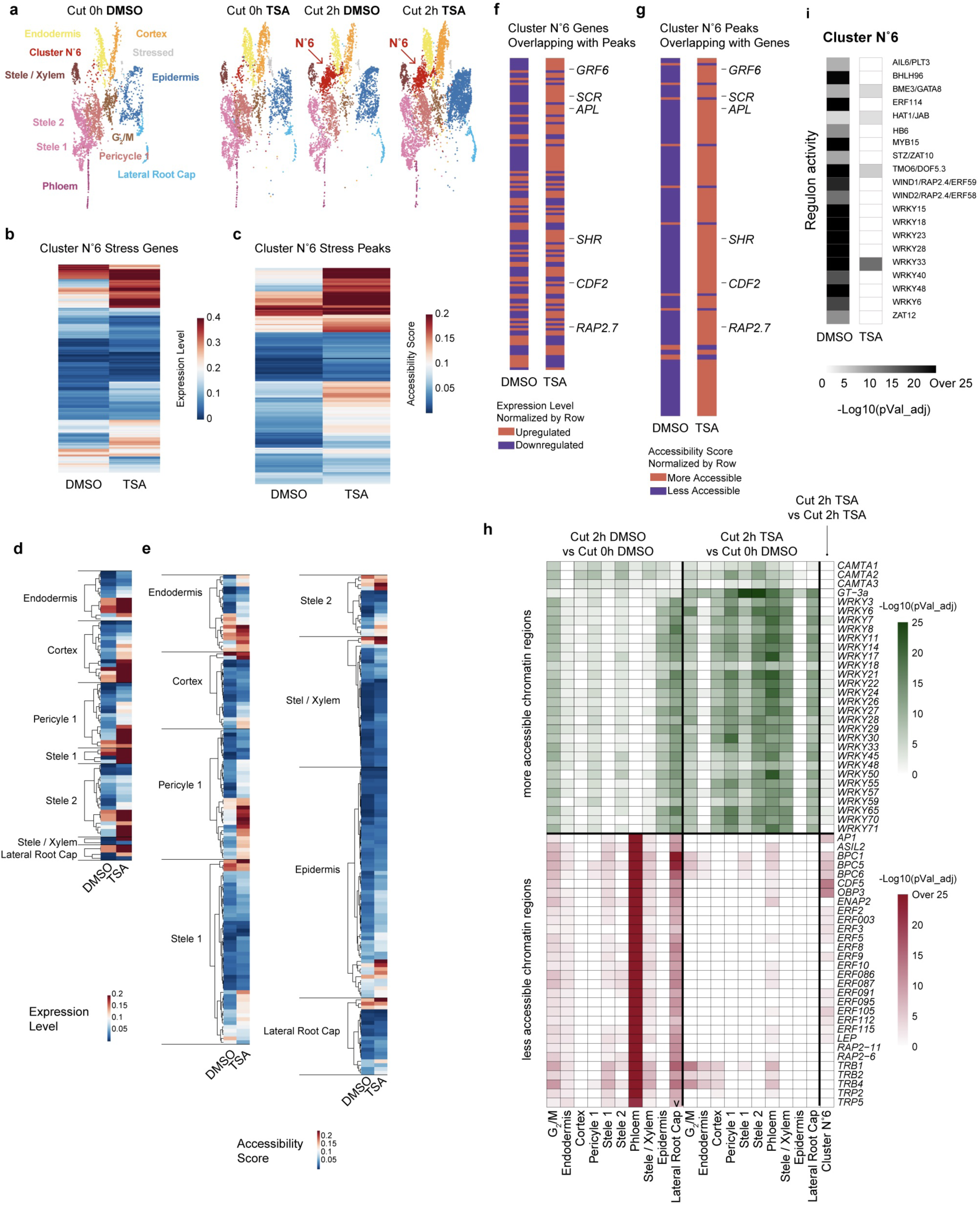
Multiome analysis reveals rapid HDAC suppression of stress and cell identity. (a) UMAPs showing integrated and separately viewed profiles of control and treated Multiome analyses. Arrow pointing to cluster N°6 shows cells unique to regeneration. (b,c) Heatmap of stress-annotated genes expression (b) or accessibility (c) showing a failure in TSA treatment to be (b) downregulated or (c) less accessible during regeneration in cluster N°6 compared to other cell types. (d,e) Heatmap of cell-identity genes expression (d) or accessibility (e) showing a failure in TSA treatment to be (b) downregulated or (c) less accessible in cluster N°6 compared to other cell types. (f,g) Genes that either are downregulated or are less accessible in cluster N°6 compared to other clusters at 2hrs post cut in control. Comparative heatmap shows the effect of TSA treatment on the same genes, showing many genes that fail to be downregulated or become less accessible or both. Genes at the bottom of the list have multiple peaks generating extra rows in g. Gene names show the behavior of key developmental genes. (h) Heatmap showing transcription factor motif enrichment in each cell type among the more (in green) or less (in red) accessible chromatin, between Cut 2h and Cut 0h. (i) Analysis of regulon activity assessed by MiniEx in DSMO and TSA treatment in cluster N°6 and other clusters.

To distinguish between the effects of the cut itself and of the short TSA treatment, we calculated the differentially expressed genes (DEG) (**Supplemental Table 4**) and differentially accessible chromatin regions (DACR) (**Supplemental Table 4**) in mock and TSA conditions using Signac’s default parameters (see Methods). Among the DACRs, and for each cell type, we analyzed transcription factor motif enrichment in chromatin regions that are either more accessible or less accessible during regeneration of mock- or TSA-treated roots (**Supplemental Table 4**). Lastly, we predicted gene regulatory networks (GRN) using Mini-Ex ^51^ and Inferelator 3.0 ^52^ to identify active transcription factors and target modules specific to each condition and cell type (**Supplemental Table 4**).

In the mock conditions, when comparing cut vs uncut roots for each cell type, we found 354 more-accessible chromatin regions and 1,571 upregulated genes in cut roots, with both gene lists enriched for stress response, auxin polar transport, and glutathione signaling as a primary response to wounding (**Supplemental Table 5**). This is consistent with independent data as wound response, auxin signaling, and reactive oxygen species signaling are all well characterized responses to injury ^53–56^.

The genes that were transcriptionally downregulated and/or became less accessible at the chromatin level in uncut vs. cut roots provided insights into the functions that need to be down-regulated in the first steps of regeneration. Genes in either category that fail to be downregulated and/or to become less accessible in TSA treatment were of particular interest, as they represent potential HDAC-regulated genes within the two-hour window. However, we also examined genes that were affected by TSA at either the chromatin or gene expression expression levels as the absence of one level of regulation could be due to high false negative rate in detecting chromatin accessibility, or the transcriptional effects of chromatin-level changes might become apparent at later time points.

We hence focused our analysis on the reprogramming cells (cluster N°6), asking what gene expression and chromatin-level changes were unique to the cluster in the two hour mock treatment, focusing on repression during regeneration as a potential direct role of HDACs. At the gene expression level, we identified 1,643 downregulated genes in cluster N°6, enriched for multiple metabolic processes but particularly stress responses, suggesting a suppression of the stress response in this reprogramming cell type compared to non-reprogramming cells (**Supplemental Figure 6g, Supplemental Table 5**). Among these genes, 70% (1,152) failed to be downregulated in the TSA 2 hour cut dataset, especially genes associated with stress response suggesting that TSA is preventing downregulation of stress responses and metabolism (**Figure 6b, Supplemental Table 5**). Similarly, at the chromatin level, we found 1,154 peaks that become less accessible in the two-hour time point in mock but stay accessible in TSA treatment (**Supplemental Figure 6h**). The closest genes associated with those peaks were also enriched for GO terms related with response to chemicals, as well as cell differentiation and root developmental process (**Figure 6c, Supplemental Table 5**). This suggests that HDACs are needed within the first two hours of regeneration to immediately downregulate the stress response instigated by wounding.

Interestingly, downregulated genes as well as less accessible chromatin regions in cluster N°6 were both enriched for cell type-specific marker genes (4 cell types) or peaks (8 cell types) respectively, but remained more expressed or accessible in the TSA treatment (**Figure 6d,e, Fisher exact test in Supplemental Table 5**). This suggests that in addition to stress repression, the early loss of cell identity during the first 2 hours of regeneration, could be mediated by HDACs. In this case, there were 145 genes that overlap between the failure to close chromatin and the failure to repress gene expression in TSA treatment vs. control (Figure 6f,g,). This overlap in the failure to downregulate gene expression and make chromatin less accessible in TSA treatment during regeneration included well-known functional root cell identity genes, such as *SCARECROW*, *SHORT ROOT*, *CLE25*, *ALTERED PHLOEM DEVELOPMENT*, *GROWTH-REGULATING FACTOR 6*, *CYCLING DOF FACTOR 2*, and *RELATED TO AP2.7* (**Figure 6f,g**), suggesting that these genes with localized roles in tissue development require chromatin closing to be downregulated during regeneration in the early stages.

This failure to repress cell identities was reminiscent of our findings with cell identity reporters that showed inhibition of HDACs by TSA led to the failure of remnant identity loss in reprogramming cells near the injured root tip (**Figure 2, 3**), even after only a 1-hour pulse of TSA (**Figure 5e**). Thus, one role of the HDACs in early regeneration is shutting down remnant cell identities that will ultimately reprogram to form the new root tip.

### WRKY and ERF modules show strong impact in early reprogramming trajectory

Looking at transcription factor motifs, we found that more accessible chromatin regions were strongly enriched for motifs of the WRKY transcription factor family in many cell types 2 hours after cut (**Figure 6h**). While these motifs were also more open in TSA in the different cell types, we found that the WRKY associated domains were opening more in DMSO compared to TSA only in the Cluster N°6 - cells in regeneration (**Figure 6h**). In parallel, *WRKY* regulons were also strongly enriched in our predicted gene regulatory networks using Mini-Ex or Inferelator in the cells in regeneration in mock treated condition but not in TSA-treated roots (**Figure 6i**, **Supplemental Table 4, Supplemental Figure 6i**). WRKYs are known to interact with OBERON (OBE) protein and this WRKY/OBE complex is known to repress stress genes and promote growth and development ^57^. We found *OBE1*, *OBE2*, *OBE3*, and *OBE4* expression to be induced during regeneration and their genomic regions to have more accessible chromatin profiles, but not in the TSA-treated condition (**Supplemental Table 4**), a possible secondary effect of blocking HDACs. These data suggest that repression of stress responses in the regeneration-specific cluster N°6 could be modulated by a WRKY/OBE module. The well-known role of WRKYs in mediating stress responses supports the data quality and suggests a specific type of stress response in reprogramming cells.

Analysis of transcription factor motif enrichments in chromatin regions that are less accessible in the different cell types after cut and in the regeneration-specific cluster N°6 cells revealed a strong enrichment for the Ethylene Response Factors (ERFs) family of genes, suggesting that chromatin associated with ERF target genes is becoming less accessible during regeneration (**Figure 6h**, **Supplemental Table 4**). We did not observe an enrichment of ERF motifs among genes that fail to become less accessible in TSA in cluster N°6. However we still see closures associated with TBRs, BPCs and ENAP1/2 that are related to methylation repression ^58–60^ showing the specificity of the TSA treatment not affecting the methylation machinery.

We found that *ERF114* and *ERF115* were specifically expressed in regeneration-specific cluster N°6 cells and that their expression was not present in TSA-treated roots (**Supplemental Figure 6j**), an apparent secondary effect of the TSA treatment. Interestingly, our MiniEx analyses showed an activity of ERF59/*WIND1 and ERF58/WIND2 and ERF114* regulons in the regeneration-specific cluster N°6, and their induction was impaired by TSA treatment (Figure 6i).

These findings suggest that the HDAC-mediated closing of chromatin regions is mediated by ERFs such as WIND1/2 or/and ERF114/115, maybe by interacting together.

### Class I HDACs promote regeneration by controlling stress

We sought to narrow down which class of HDACs mediated early regeneration responses by examining inhibitors of various HDAC classes (**Supplemental Figure 7a,b**). These experiments indicated that inhibition of class I HDACs had the strongest effect on regeneration in our system. We therefore set out to test the role of class I HDACS and, specifically, the unexpected role in suppressing stress responses early in regeneration. One tool that could help parse the importance of repressing stress responses from other roles in chromatin modification were the opposing effects of class I HDACs *HDA6* and *HDA19* in relation to stress. While *HDA19* and *HDA6* function redundantly in several contexts ^61–64^, they also are known to have opposing roles, for example, in salt stress response ^47^. *HDA6* was shown to promote the salt stress response, while *HDA19* repressed the salt stress response ^47^. If a major role of HDACs in early regeneration is repressing stress responses, as implicated by the Multiome analysis, we reasoned that *HDA6* and *HDA19*, with opposite roles in salt stress response, should show opposing mutant phenotypes in regeneration. Therefore, we tested the two loci and other Class I HDAC mutants for regeneration, using the sensitive mPS-PI amyloplast assay.

We tested two *hda19* mutant alleles: *SALK_0277241*, and *hda19-3*. Both mutants had significantly fewer amyloplast-positive cells at 18h compared to wild type (**Figure 7a**). We also tested three *hda9* mutants (*SALK_067576*, *SALK_083935*, and *SALK_007123* (*hda9-1*), with two having significantly fewer amyloplast-positive cells at 18 hours compared to wild type (**Figure 7b**). Mutants for *HDA7,* which was recently shown to lack deacetylase activity ^65^, regenerated similarly to wild type roots *(SALK_002912* and *SALK_108050,* **Figure 7c**). Overall, the results with *HDA19* and *9* are consistent with TSA treatment, suggesting that HDAC activity and chromatin closure promote root regeneration.

**Figure 7.**
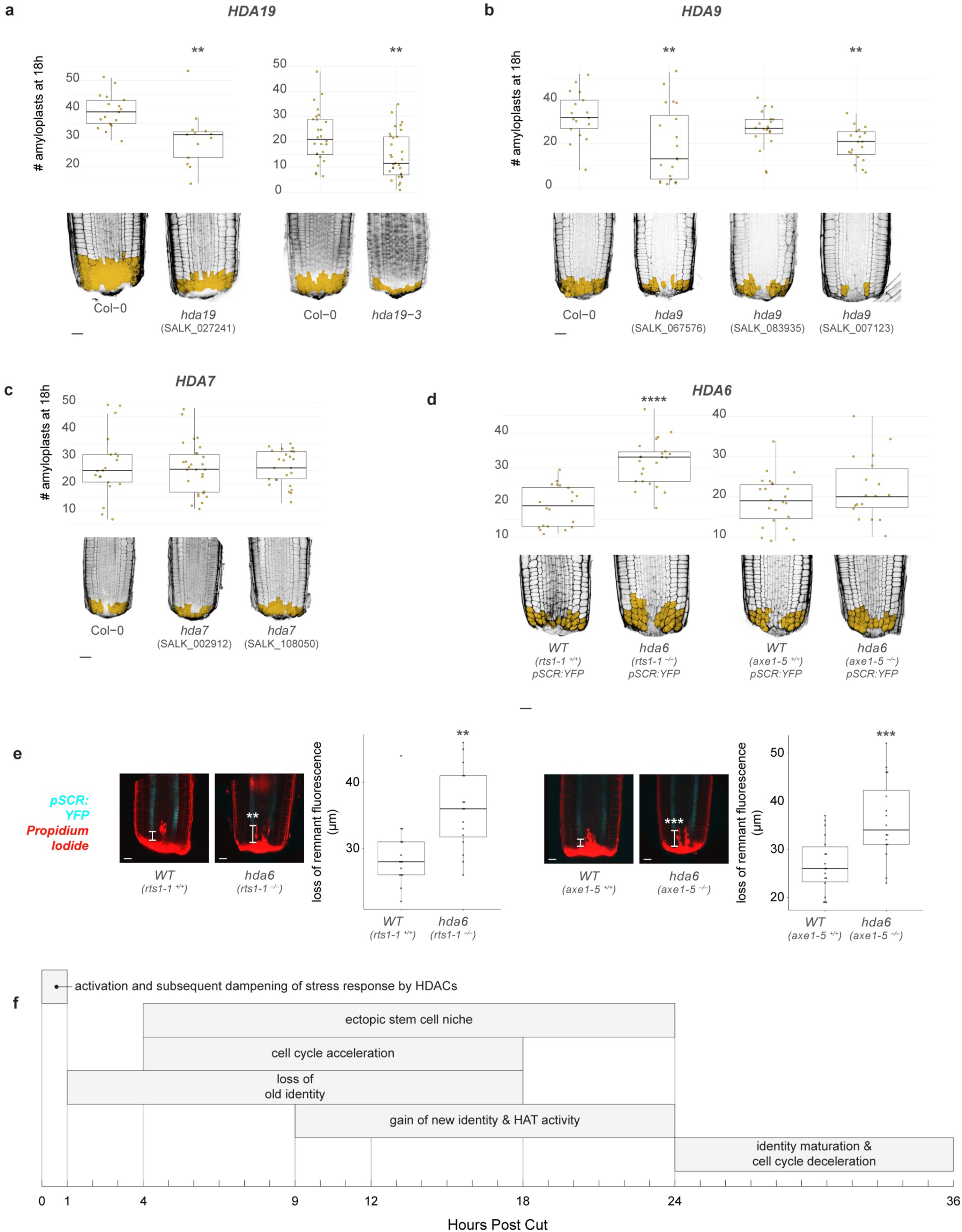
Regeneration phenotypes of Class I HDAC mutants. (a) Boxplot (top) showing number of amyloplast-positive cells at 18 hours post cut in wild type (Col-0) or *hda19* mutants as seen by mPS-PI staining (bottom). (b) Boxplot (top) showing number of amyloplast-positive cells at 18 hours post cut in wild type (Col-0) or *hda9* mutants as seen by mPS-PI staining (bottom). (c) Boxplot (top) showing number of amyloplast-positive cells at 18 hours post cut in wild type (Col-0) or *hda7* mutants as seen by mPS-PI staining (bottom). (d) Boxplot (top) showing number of amyloplast-positive cells at 18 hours post cut in wild type (Col-0) or *hda6* mutants as seen by mPS-PI staining (bottom). (d) Confocal microscopy of *hda6* mutants expressing *pSCR:YFP* with corresponding boxplots showing the loss of remnant fluorescence at the cut site. Asterisks indicate significance after t-test: ** = p<0.001, *** = p<0.0001. (f) Temporal map of the major regeneration events outlined in the present study. Scale bars in (a)–(e) correspond to 20 µm.

On the other hand, two *hda6* mutant alleles, *rts1-1* and *axe1-5* both had more amyloplasts at 18 hours, although only *rts1-1* was significant (**Figure 7d**), suggesting improved regeneration compared to wild type. We crossed the *rts1-1* and *axe1-5* mutant lines into a transcriptional reporter for *SCR*, and measured the amount of remnant identity loss in mutants vs wild type. Consistent with the mPS-PI results, both *rts1-1* and *axe1-5* mutants had a larger region of diminished *SCR* expression near the wound site, suggesting a more rapid loss of remnant cell identity (**Figure 7e**). Taken together, these results suggest that *HDA6* has a negative role in regeneration, with defects in suppressing the stress response associated with wounding. The opposing roles of *HDA9/19* and *HDA6* in stress and their opposing regeneration phenotypes are consistent with a requirement to down-regulate stress during regeneration.

Because Class I-specific inhibition did not have a total inhibitory effect on regeneration to the extent that TSA did (**Supplemental Figure 7c-f**), we also tested mutants for *SRT* and *HD-tuin* family HDACs (**Supplemental Figure 7g,h**). We found that one *HD2* family HDAC—*HD2D*—had significantly fewer amyloplasts at 18 hours compared to wild type (**Supplemental Figure 7h**), suggesting poorer reprogramming efficiency. Across the different cell types, we found 130 chromatin regions that became less accessible in the 2 hour cut condition after injury, and these regions were enriched for targets of the related *HD2B* ^66^ suggesting a role for the plant-specific HD-tuin HDACs in this process (**Supplemental Table 5**).

Taken together, the data is supportive of a model in which one early role of a subset of Class I HDACs is to rapidly downregulate a stress response instigated by injury but detrimental to regeneration. Another role of the HDACs appears to be shutting down remnant identities to allow new cell identities to specify and proceed with cellular reprogramming.

## Discussion

We found that HDACs act within the first hour of wounding to license efficient reprogramming by dampening a runaway stress response and mediating the loss of remnant identities. From there, meristematic youth is overlaid and cells rapidly change identities to replenish missing cell types removed after injury. We implicated *HDA9* and *HDA19* as positive regulators of regeneration, and *HDA6* as a negative regulator. This is correlated with their known roles in stress responses—*HDA9* is a negative regulator of salt and drought stresses ^67^, whereas *HDA6* is a positive regulator of salt stress responses ^7,68^. This supports conclusions from the Multiome analysis and TSA treatments that one of the primary roles of HDACs in the first hours after injury is to suppress a runaway stress response that was instigated by wounding. Simultaneously, HDACs mediate the loss of remnant identities, a process that results only in a brief period of overlapping cell identities. Thus, while wounding is clearly necessary to induce regeneration, the plant must actively contain at least some aspects of the wounding response in order to successfully regenerate. It is also possible that in other scenarios, such as where wounding is part of an ongoing attack from a soil pest or pathogen, the full wounding response is advantageous. In addition, the data shows that, in the context of root tip regeneration, the loss of cell identity is not just a lack of instructive cues resulting from a new signaling environment but, rather, the plant actively and rapidly erases remnant identities to make way for new ones.

During normal growth, cell division and differentiation are coupled, along with growth itself ^69^. This makes experimental approaches to understanding any one of these processes particularly challenging, as perturbations of one will necessarily influence the rest. Root tip regeneration, as an experimental system, circumvents this issue by resetting development in an adult context. By forcing cells to reprogram after having already been specified, the timings of division and identity establishment can be unlinked. Still, given the high background mitotic activity in the root meristem, and the ongoing specification of cells prior to the differentiation zone, pharmacological and single-cell multiomics approaches were used to further tease apart division and differentiation processes.

The results show that, during regeneration, cell division is necessary to respecify cell identities that were removed by cutting. Interestingly, genes implicated in early root specification, such as *WOX5* and *WIP4,* show a rapid return that is division-independent. Thus, expression of some of the genes that characterize the re-establishment of the meristem are division-independent, particularly those expressed early and most directly responsive to wounding. This shows that cells in the root tip do not need cell division to embark on respecification, but they do need division to make the full transition to a new identity. What aspect of cell division, such as replication-dilution models or other mechanisms, such as phase-specific expression of chromatin remodelers, remains an open question.

Additionally, the cell cycle is accelerated nearly 3-fold compared to cells in the same region in uncut roots. Acceleration of the cell cycle is commonly observed in contexts where new tissues are being differentiated, such as lateral root formation and embryogenesis in plants ^70,71^ , or gastrulation in rats ^72^. This acceleration is typically achieved by shortening G_1_ and/or G_2_. Indeed, rapid cell cycles with short G phases are correlated with high programming rates in induced pluripotent stem cells ^73^. In this work we suggest that the plant finely tunes division rates during regeneration, with first an *OBP1*-mediated acceleration to facilitate reprogramming, then an *SMR*-mediated slowing down to enable maturation. We also showed recently that a glutathione burst in the region of regenerating cells within seconds after wounding promotes shortened G1, rapid cell divisions, and cellular reprogramming ^19^. Thus, wounding itself triggers rapid cell cycles to speed the reprogramming process but it appears necessary to subsequently put the breaks on cell division, as we show through the *SMR* mutant analysis. *SMR5* was among the genes we found specifically induced at the transcriptional level during regeneration. Thus, both acceleration and deceleration mechanisms appear to be wound-responsive, albeit with a delay in deceleration. Overall, this shows how the plant finely tunes the necessary division rates in regeneration to reprogram cell identity.

In contrast to the significant influence of HDACs on regeneration, we found a relatively minor contribution of HATs, mostly later in regeneration. Interestingly, Rymen et al^46^ found that inhibition of the GNAT-MYST family of HATs, whether by pharmacological treatment with MB-3 or mutation of *HAG1* or *HAG3*, led to reduced callus formation, whereas inhibition of the CBP family of HATs via C646 treatment did not ^46^. We, on the other hand, found the opposite effect in our root tip regeneration system: MB-3 had no effect whereas inhibition of CBP HATs with Curcumin treatment had a stronger effect. This could point to different roles for different classes of HATs in different regeneration contexts.

Our chemical treatments allow us to create a temporal map of the processes needed at different stages of regeneration (**Figure 7f**). Within the first two hours, the closure of chromatin, mediated by Class I HDACs, suppresses an initial burst of stress response genes and facilitates the repression of remnant identities. Next, starting at 4 to 9 hours, a gene expression program associated with the stem cell niche and the young meristem activates broadly and ectopically, concomitant with an acceleration of the cell cycle to three-fold its homeostatic rate driven in part by *OBP1*. These early genes, many of which have specific roles in stereotypical root development, appear to be a part of the stress response that are independent of cell division or the remodeling of chromatin acetylation. The final components of cell identity appear to depend on cell division, but the rapid divisions instigated by injury ensure this process occurs quickly, with our time course of single-cell genome-wide analysis showing that markers for specific cell identities begin to appear around 14 hours after injury. This process appears ongoing at the transcriptional level with loss and gain proceeding somewhat in tandem. CBP family HATs act at the latter end of this stage to promote new identity formation. Finally, the cell cycle slows down, mediated in part by *SMR5*, *SMR7*, and *SMR10,* to transition to mature cell types as the regeneration process is completed. This analysis illustrates how the plant actively manages fundamental processes to complete the trajectory from remnant identity to new cell identities within 36 hours in the root tip.

## Methods

### Plant Growth and Maintenance

Seeds of Arabidopsis plants were stratified at 4°C for 2 days, then sterilized and placed on agar plates containing ½ Murashige and Skoog salts (Sigma M5524), 0.5% sucrose, and grown vertically in chambers set to 23°C and a 16h light / 8h dark cycle (80-90 μmol m^-2^ s^-1^) for one week.

### Microsurgery

Removal of root tips was done by our previously published microsurgery protocol ^17^. Briefly, excisions were done roughly 130 µm from the tip of the root, by hand, with a 30G sterile dental needle (ExcelInt) or just above the vascular initials by using a microscalpel (Feather 72045-15) under a dissecting microscope with Rottermann Contrast to visualize the upper limit of amyloplasts within the root tip.

### Live Imaging

Live imaging of regenerating roots was done using a modified version of our previously published protocol for tracking long-term growth in roots ^40^. As the QC and therefore endogenous *WOX5* domain is removed for regeneration studies, the fluorescent region marked by pWOX5:GFP used for tracking is lost. As a workaround, I tested several live dyes with emission profiles outside that of our cell and tissue reporters.

A variety of dyes, including DAPI and the SYTO family of nucleotide labels, were tested. Ultimately the best solution was found to be DiD’ (Life Technologies), a lipophilic, non-toxic live dye designed to adhere to plasma membranes, have slow diffusivity, and persist for several days while retaining bright fluorescence. DiD’ brightly labeled the cut root tip in several clusters, which gradually shrink to discrete, cell-sized punctae that are amenable to tracking. As these punctae can appear at various positions along the root’s radial axis, minor drift can occur during extended periods of acquisition as the drift correction module selects different reference planes. As such, while the uncut root can effectively be imaged in a fully automated manner, tracking regenerating roots requires frequent manual adjustment to settings or to the microscope stage itself (see detailed Materials/Methods). The first few hours, in particular, are critical and require active on-the-fly adjustment of the microscope controls to ensure successful imaging.

In lieu of the usual QC-localized fluorescent reporter, a fluorescent DiD’ cluster can be presented as a single, discrete object for MatrixScreener to track. Once a suitable cluster is found. To localize dyes to the regenerating root tip, we used a modified version of our microsurgery protocol (Sena et al 2009). By dipping the needle in a solution of DiD’ before using it to cut away the root tip, the dye can be primarily localized to the cut site.

### Pharmacological Treatment

For experiments with cell cycle inhibition (Roscovitine), chromatin modifier inhibition (Trichostatin A, Circumin, and MB-3), and EdU, the chemical in question was added to autoclaved liquid MS media that had been cooled to 50°C. Seedlings were transferred to plates containing inhibitors immediately before root tip removal.

### EdU and f-ara-EdU Labeling

A modified version of Kazda et al’s protocol ^74^ was used for EdU labeling. Rather than keeping seedlings in the same dish and replacing the solution at each step, a small dedicated petri dish was used for each solution. That is, seedlings were transferred from a petri containing fixative, to one containing wash buffer, and so on. Rather than use a commercial click cocktail solution, we used a formulation based on a protocol described <u>on ResearchGate</u>. The labeling solution contained the following reagents at the stated final concentrations in PBS: 2 mM Copper(II) Sulfate Pentahydrate (Sigma 12849), 8 µM Sulfo-Cyanine5 Azide (Lumiprobe B1330), and 20 mg/mL L-Ascorbic Acid (Sigma A4544), added to PBS in that order. DAPI counterstaining was done as described by Kazda et al.

f-ara-EdU labeling to assess mitotic activity in the *obp1* and *SMR* mutants was adapted from Goldy et al 2021 ^75^. Wild type and mutant plants were either treated with a 15 minute pulse of f-ara-EdU followed by a 4 or 8 hour chase, or kept on f-ara-EdU for 3, 6, 9, or 12 hours. In the pulse-chase experiment, a median confocal image of the root was taken, and within the meristematic zone (from cut tip up to the first doubled cortical cell) the proportion of mitotic figures over total EdU-positive nuclei was counted. In the experiments where plants were kept on f-ara-EdU for 3, 6, 9, or 12 hours, a confocal image of the epidermis was taken, and within the meristematic zone (from cut tip up to the first doubled epidermal cell) the proportion of EdU-positive nuclei over total (DAPI-positive) nuclei was taken.

### mPS-PI staining

mPS-PI staining was done following Truernit et al. ^44^. Briefly, seedlings are fixed in 10% acetic acid, 50% methanol for at least one hour, or up to several days. Fixed seedlings are rinsed with water, incubated in periodic acid for 1 hour, rinsed again, incubated in Schiff reagent with propidium iodide for 2-3 hours, then incubated overnight in Visikol for Plant Biology as a clearing agent. Cleared seedlings were mounted in either Visikol® for Plant Biology™, Image-iT™ Plant Tissue Clearing Reagent, or ClearSee, and imaged using a confocal microscope with 561 nm excitation wavelength.

### Lugol staining

Seedlings were fixed in 50% ethanol solution at 48 hours post cut, and stored overnight at 4° C or up to several weeks. After acclimating to room temperature, seedlings are briefly swirled (five circular motions) in iodine, lugols, 5%, and transferred to a glass slide with a small volume of clearing agent (Visikol® for Plant Biology™, Image-iT™ Plant Tissue Clearing Reagent, or ClearSee) applied beforehand. After mounting, a few more drops of clearing agent are added to the slide to fully clear what is left of the lugol. Slides were imaged under a stereo dissecting microscope with no coverslip and scored for regeneration.

### Mutant Lines

To generate the CRISPR SMRs SMR5 (AT1G07500), SMR7 (AT3G27630), SMR10 (AT2G28870) lines we used the PYUU CRISPR plasmid (https://www.addgene.org/159751/) from Angulo et al ^76^. gRNA were designed using the pipeline Chopchop (https://chopchop.cbu.uib.no/ ) with default parameters. The cassettes Prom:gRNA:scaffold:Terminators were synthesized and inserted into PYUU using the goldengate method. One cassette contained three gRNA against SMR10, another cassette contained four gRNA, two against SMR7 and two against SMR10. A third cassette contained two gRNA against SMR10, one against SMR5 and one against SMR7 (**Supplementary Table 6**). To obtain transgenic lines, the PYUU plasmids were transformed into the Agrobacterium strain GV3101, which were then transformed into wild-type plants of Arabidopsis Col-0 using the floral dip method. T2 and T3 lines were sequenced around the gRNA site to identify mutated homozygous plants.

T-DNA mutants were ordered from ABRC and genotypes confirmed by PCR: AT3G50410 *obp1* (SALK_049540), AT5G55760 *srt1* (SALK_064336C), *srt1-1* (SALK_086287C), AT5G09230 *srt2* (SALK_131994C), *srt2-2* (SALK_149295C), AT5G22650 *hd2b* (SALK_113502), AT5G03740 *hd2c* (SALK_136925C), AT2G27840 *hd2d* (SALK_095273), AT4G38130 *hda19* (SALK_027241), hda19-3 (SALK_139445), AT3G44680 *hda9* (SALK_067576), *hda9* (SALK_083935), *hda9* (SALK_007123), AT5G35600 *hda7* (SALK_002912), *hda7* (SALK_108050), AT5G63110 *hda6 (rts1-1)*, *hda6 (axe1-5)*.

### Cell and nuclei generation

Protoplast isolation was done as previously described ^77^, with slight modifications. Protoplast solution is made fresh on day of protoplasting: 400 mM/8% mannitol, 20 mM MES hydrate, 20 mM KCl, 20 mM CaCl2, 1.2% Cellulase R10 (Yakult,ONOZUKA R-10, Japan), 0.4% Macerozyme R10 (Yakult, Japan) in 10 mL water. The solution was adjusted to pH to 5.7 with 1 M Tris pH 8. Add 0.1% BSA and filter using a 0.4um filter. About 300 roots are transferred in 2 ml of protoplast solution in a small petri dish (35×10mm) for 15mins at 100 rpm. As the enzyme degrades preferentially the elongation zone, when the root tips are detached, they are transferred into a new small petri dish containing 2 ml of fresh protoplast solution. Root tips are incubated for 1h45 at 100 rpm at room temperature. After passing through two 20um filters placed on top of a 40 μm strainer, protoplasts were centrifuged at 500 g for 5 min and resuspended twice in washing buffer: 400 mM/8% mannitol, 20 mM MES hydrate, 10 mM KCl, 10 mM CaCl2, pH 5.7, BSA 0.1%, filtered at 0.22um. Final suspension volume was adjusted to a density of 2,000 cells/μl. Protoplasts were placed on ice until further processing.

For multiome, roots were grown on MS plate as described above but on a nylon mesh (110um). The mesh was then transferent MS plates with DMSO or TSA (10uM) and cut immediately at 130 µm from the tip of the root. Alternatively, control roots were transferred on DMSO or TSA and kept for 2h without cutting. After 2h, roots are cut again 100um above the first cut and picked with a pipette tip containing the nuclei lysis buffer. Control roots (defined as cut0h) were then cut 130 µm from the tip of the root and immediately picked with a pipette tip containing the nuclei lysis buffer to be processed. About 150-200 root tips were processed per sample on the same day to be processed in parallel. Nuclei were extracted by mechanical tissue disruption, and centrifugation as described in Guillotin et al. 2023 ^78^, but nuclei were resuspended in 10xGenomics nuclei buffer at the end instead of the Final buffer from our protocol.

### Single Cell Sequencing

16,000 cells per replicate were loaded in a Single Cell B, and G Chip (10x Genomics). Single-cell libraries were then prepared using the Chromium Single Cell 3’ library kit V3 and V3.1, following manufacturer instructions. Multiome was performed using 10x genomics Chromium Next GEM Single Cell Multiome ATAC + Gene Expression kit following the manufacturer protocol. Libraries were sequenced with an Illumina Novaseq 6000 chip SP V2.5 (4 libraries per chip). Raw scRNA-seq data was analyzed by Cell Ranger 5.0.1 (10x Genomics) to generate gene-cell matrices. Gene reads were aligned to the Arabidopsis TAIR10.38 genome.

### Single Cell Analysis

Sequenced cell replicates (see Supplementary Table 1) were integrated and cells mapped using the Seurat package v4.1 as follows: first, genes with counts in fewer than three cells were excluded from the analysis and their counts were removed. Second, low-quality cells having less than 1600 genes and 10 000 UMI or more than 11 000 genes or 400 000 UMI detected were removed from the analysis as bad quality cells and potential cell doublet. Clustering of cells separately were done using the seurat ’SCT’ approach ^35^. First raw reads were normalized using the SCTransform function, then SelectIntegrationFeatures was used to identify anchors between the datasets, using 3000 features. Control replicates were used as reference datasets to map the cut replicates. Finally, a Principal Component analysis (PCA) is performed using the first 50 principal components and a non-linear dimensional reduction was performed using the UMAP algorithm with the top 50 PCs.

### Topic Modeling

The number of suitable topics has been established using the R package dynRB to empirically identify how many topics are necessary to distinguish every cell type in the control condition with a minimum of overlap between them. Eleven topics were found to be the minimum number with the smallest overlap compared to K+1, whereas any more than 11K appears to overfit the model. The R package CountClust (https://github.com/kkdey/CountClust) was then run on the single cell count expression matrix, containing all the cells from ctr and cut dataset without prior labeling. Gene characteristics of each Topic were identified using the ExtractTopFeatures function from CountClust. Cells having a proportion of orange topic above 0.21 were selected as “in regeneration”

### Mini-Ex Analysis

We used the package MiniEx ^51^. as described in https://github.com/VIB-PSB/MINI-EX on the single cell expression count matrices for each cell type and each time point.

### Identification of genes involved in root regeneration

To identify the genes involved in cells in regeneration we combined several statistical analyses:

- To identify the genes that have their expression modified during the regeneration process, we performed differential gene expression analysis using the FindMarker function from Seurat using default parameters but with min.pc= 0.01, pval < 0.05. The test was conducted in the cells in regeneration found by topic modeling and between the different time points (eg: cells in regeneration Cut 4h vs cells in regeneration Cut 9h etc..).
- To improve this prediction we also performed differential gene expression analysis using the same parameters, for each cell type between the cut dataset and the respective controls (eg: Cell type 1 in Cut 4h vs Cell type 1 Ctr 4h etc..).
- To identify the genes that were enriched in cells in regeneration vs the other cell type we also performed a differential gene expression analysis using the FindMarker function from Seurat using default parameters but with min.pc= 0.01, pval < 0.05. This time we tested between the cells in regeneration vs all the other cells in the same biological replicate (eg: cells in regeneration Cut 4h vs all the other cells in Cut4h).
- Finally to only look at genes that are only expressed in specific cell type and not in every cell type we added the tau value of each genes (Yanai et al., 2005^)^ 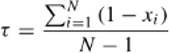 where N is the total number of cell types and xi is the expression profile component normalized by the maximal component value. We only considered genes with a tau of at least 0.7 in Ctr or Cut datasets.

The compiled dataset is displayed in Supplementary Table 3 NB: the FindMarker function from Seurat uses a bonferroni correction based on the total number of genes in the dataset, which is very stringent to remove actual differentially expressed genes, when expression is low or for lowly expressed genes. As here we sometimes use small cell subset in regeneration this could lead to missing numerous important genes. Hence to avoid this bias we calculated all the DEG for each cell type between the merged control and cut conditions (pval < 0.05) and removed from our analyses, all the genes found as DEG in 75% or more of the cell type, and that we consider as batch effect.

### Identification of genes affected by chemical treatments

To identify genes affected by TSA, roscovitine and curcumin, we integrated them separately with our ctr/cut datasets. Indeed common integrations of the three treatments together was affecting the clustering of the control dataset itself. We created 3 seurat objects each containing ctr4h, cut4h, ctr9-14h, ctr24h, cut9h, cut14h, cut24h, DMSO14h and one treatment (TSA or roscovitine or curcumin) 14h. We selected 6 clusters where the cell in regeneration identified by countclust were located in these 3 different integrations. We then performed differential gene expression analysis for each of these clusters using the FindMarker function from Seurat using default parameters but with min.pc= 0.01, pval < 0.05, between the treatment and the DMSO control to identify genes affected or not by the treatments.

Supplementary Table 3 shows the comparison of the different DEG for each treatment.

### GO term analysis

All GO enrichments were performed using shinyGO V0.82 (http://bio-informatics.sdstate.edu/go/) with an FDA of 0.05.

### Gene Regulatory Network Inference with Inferelator 3.0

We used Inferelator 3.0 to infer gene regulatory networks using our single-cell gene expression data and previously published TF-DNA binding information from DAP-seq (O’Malley, 2017; Gibbs, 2022). In brief, Inferelator 3.0 first estimates a TF’s activity from the expression of its putative target genes. It then employs regularized linear models to regress target gene expression from the expression of its TF regulators, which is weighted by TF activity.

Using our previously identified regenerating cell cluster at the 2-hour timepoint, we calculated transcription factor activity using default parameters and generated the regulatory network model using Bayesian Best Subset Regression. This was done for DMSO- and TSA-treated cells separately. The algorithm was run with a total of five bootstraps, and a TF-target edge was assigned a confidence measure that corresponds to the frequency with which the edge appears across each bootstrap, e.g., if an edge appears in 3 of 5 bootstraps, it is given a confidence of 0.6. 20% of the DAP-seq binding data was left-out of training and used for model performance evaluation using Matthews Correlation Coefficient (MCC).

### Mulitome analysis

Multiome data was integrated using the R package Signac 1.8. Both chromatin accessibility and gene expression were used for clustering. All scripts used are publically available on our Github repository (see Data availability). Briefly, nuclei with genes with counts in fewer than three nuclei were excluded from the analysis and their counts were removed. Second, low-quality nuclei having less than 400 genes and 500 UMI or 500 peaks or more than 11 000 genes or 400 000 UMI detected were removed from the analysis as bad quality nuclei and potential cell doublet. Clustering of nuclei was done using the seurat ’SCT’ approach (Hafemeister, C. & Satija, R. Normalization and variance stabilization of single-cell RNA-seq data using regularized negative binomial regression. Genome Biol. 20, 296 (2019).). First raw reads were normalized using the SCTransform function, then SelectIntegrationFeatures was used to identify anchors between the datasets, using 3000 features. Principal Component analysis (PCA) is performed using the first 50 principal components and a non-linear dimensional reduction was performed using the UMAP algorithm with the top 50 PCs. Then ATACseq data was integrated and best peak features were found using the FindTopFeatures (min.cutoff = 5), RunTFIDF and RunSVD. RNA and ATAC were then integrated together using the FindMultiModalNeighbors function and a UMAP was generated. Because we did not remove the nuclei having high plastid reads, we obtained 2 clusters in the umap having very low cell markers, and mostly plastid markers. These two clusters were removed from the analysis as they were composed by low quality cells.

Then Peak calling was refined using the MACS2 package and using the Narrow parameter and the calling done on a PerCell type basis instead of the entire dataset.Differentially accessible chromatin regions between two cell types/conditions were calculated using the FindMarker function from Signac using the logistic regression ’LR’ parameter on the latent.vars being the peak called by MACS2.Differential gene expression analysis using the FindMarker function from Seurat using default parameters.Transcription factor motif enrichment was performed using the FindMotifs function from Signac, defining first the background motif and using JASPAR2022 as motif database.

### Re-analysis of ATACseq

ATACseq data from Hernández-Coronado et al., 2020 was used for this analysis. Briefly, in this paper, cells expressing Monopteros (ARF5) after cut were FACS sorted and ATACseq was performed compared to uncut roots. Open chromatin regions were then identified. Here, the percentage of genes with differential ATACseq peaks associated with gene expression (atacpct) was calculated per cell by counting all genes in each cell that (1) carry differential ATAC peaks in the promoter region and (2) are expressed greater than 1 in the sc-RNA-seq normalised dataset. That number was then divided by the total number of total differential ATACseq peaks detected and represented as a percentage. Values were then visualized on a per cell basis in UMAP space where cells were colored based on the atacpct.

### Data availability

Reference genomes were downloaded from Arabidopsis TAIR10.38, at https://www.arabidopsis.org/. All raw single-cell and single-nucleus RNA-seq data, expression matrices and analysed R-Seurat objects are available under Gene Expression Omnibus accession GSXXXX. Script used to integrate the different datasets and to perform major analyses are compiled and available on GitHub (https://github.com/BrunoGuillotin/HDACs-repress-runaway-stress-and-cell-identity-to-promote-reprogram ming-in-root-regeneration).

## Acknowledgements

This work was funded by the National Institutes of Health (R35GM136362) to K.D.B and Human Frontiers Science Program (LT000972/2018-L) to B.G. We also acknowledge the Zegar Family Foundation for their generous support.

---

**Supplemental Figure 1.**
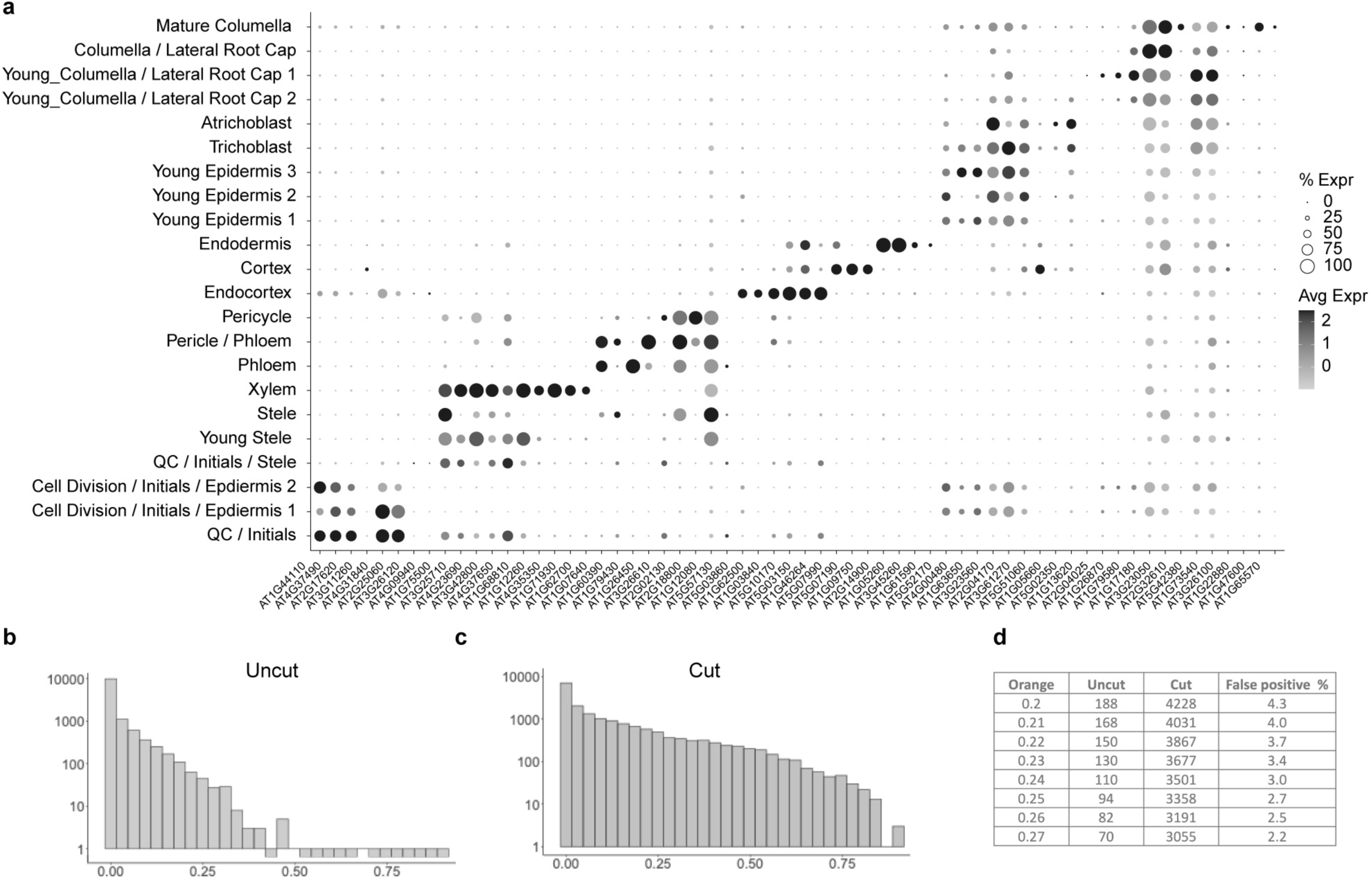
Identifying early stage regeneration markers. (a) Dotplot showing cluster-specific gene expression from scRNA-seq data in control uncut condition. (b,c) histograms showing the number of cells (y) and their proportion of orange topic predicted in our Countclust analysis, in the control uncut condition (b) or cut conditions (c). (d) contingency table of cells having orange topic proportion between uncut and cut dataset.

**Supplemental Figure 2.**
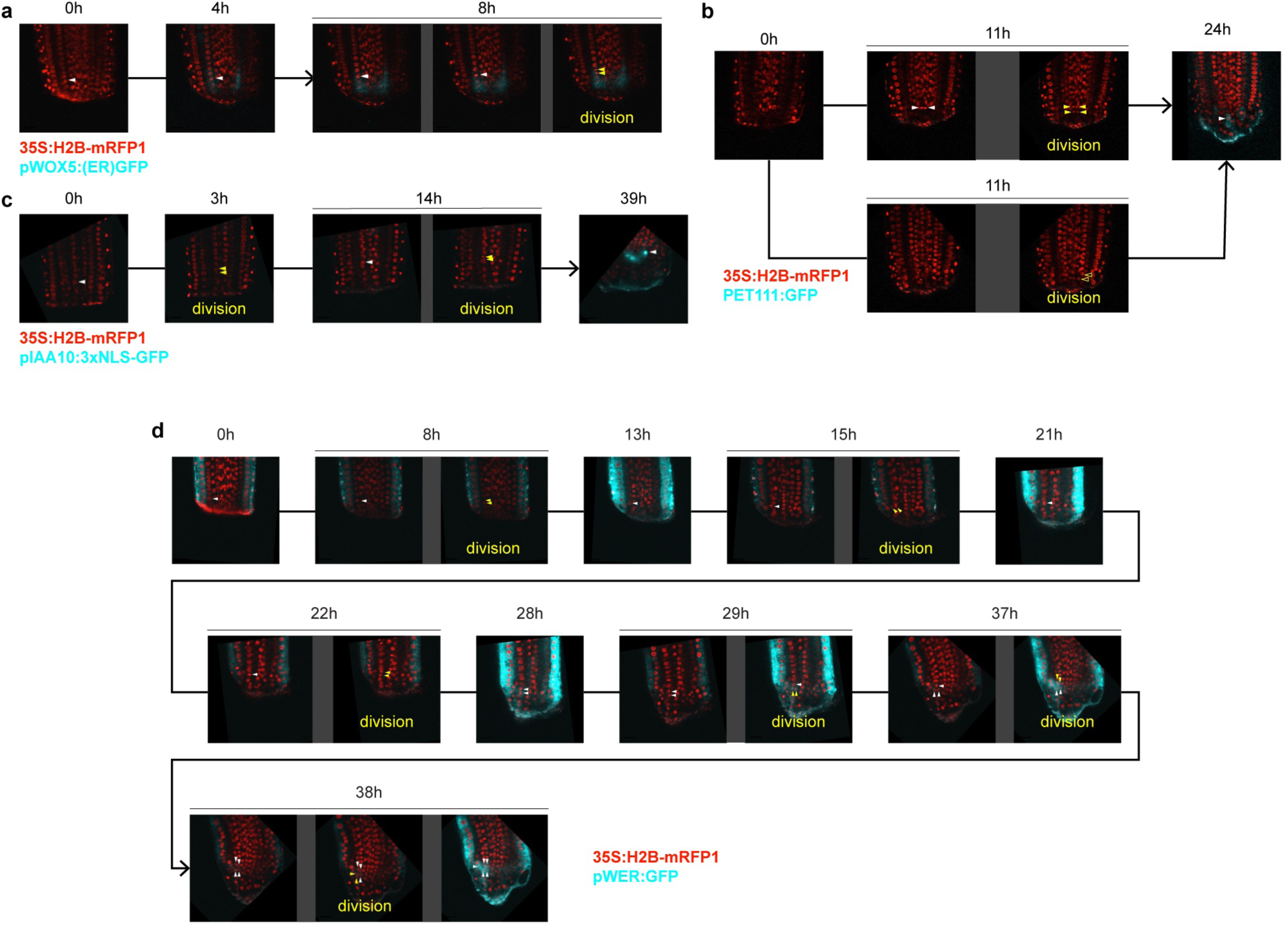
The reappearance of different cell identity markers have varied dependencies on cell division. (a) Still images from a time-lapse movie showing *WOX5* expression returning before cell division in a regenerating root. (b) Still images from a time-lapse movie showing the return of PET111 expression during regeneration occurring after cells divide once. (c) Still images from time-lapse movie showing *IAA10* expression during regeneration returning only after cells have divided twice. (d) Still images from time-lapse movie showing *WEREWOLF* expression during regeneration returning to the lateral root cap only after cells have already divided several times. Scale bars correspond to 20 µm.

**Supplemental Figure 3.**
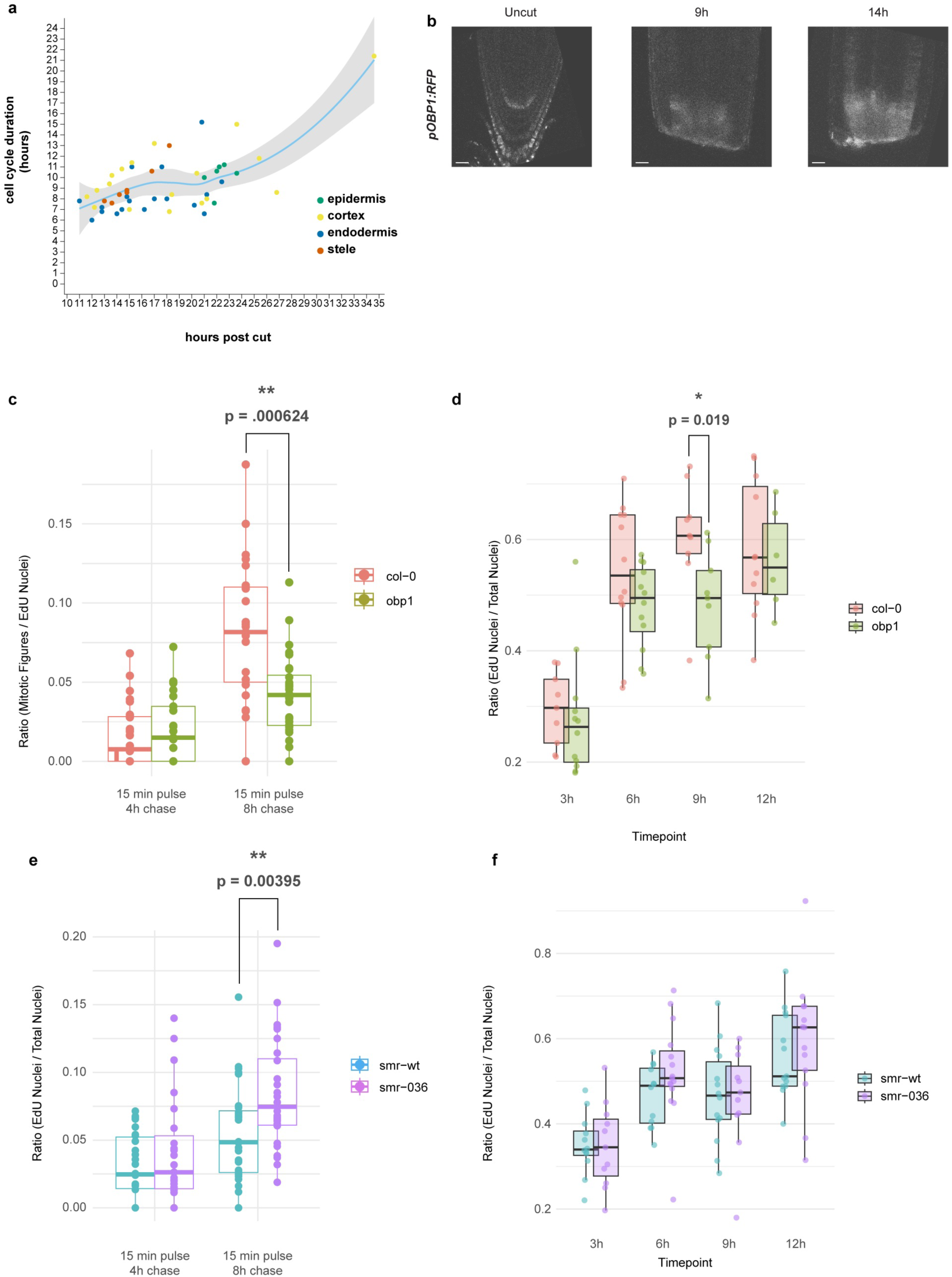
Characterization of mitotic activity in wild type and cell cycle mutants. (a) Scatterplot showing cell cycle duration for different cells during regeneration, as observed in time-lapse movies. Compared to uncut roots, the average cell cycle duration is nearly three-fold faster. (b) Confocal images of *pOBP1:RFP* expression in uncut, 9, and 14 hour post cut roots showing broad activation of *OBP1* at both cut time points. Scale bars correspond to 20 µm. (c) Boxplot quantifying ratio of f-ara-EdU-positive mitotic figures relative to f-ara-EdU-positive nuclei after a 15 minute pulse of f-ara-EdU followed by either 4 or 8 hours chase in regenerating wild type and *obp1* mutants. (d) Boxplot showing ratio of EdU-positive nuclei over total (DAPI-positive) nuclei after 3, 6, 9, or 12 hours incubation on f-ara-EdU in regenerating wild type and *obp1* mutants. (e) Boxplot quantifying ratio of f-ara-EdU-positive mitotic figures relative to f-ara-EdU-positive nuclei after a 15 minute pulse of f-ara-EdU followed by either 4 or 8 hours chase in regenerating wild type and *SMR5/7/10* triple mutants. (f) Boxplot showing ratio of EdU-positive nuclei over total (DAPI-positive) nuclei after 3, 6, 9, or 12 hours incubation on f-ara-EdU in regenerating wild type and *SMR5/7/10* triple mutants. Asterisks indicate significance after t-test: * = p < 0.05, ** = p < 0.001, *** = p < 0.0001.

**Supplemental Figure 4.**
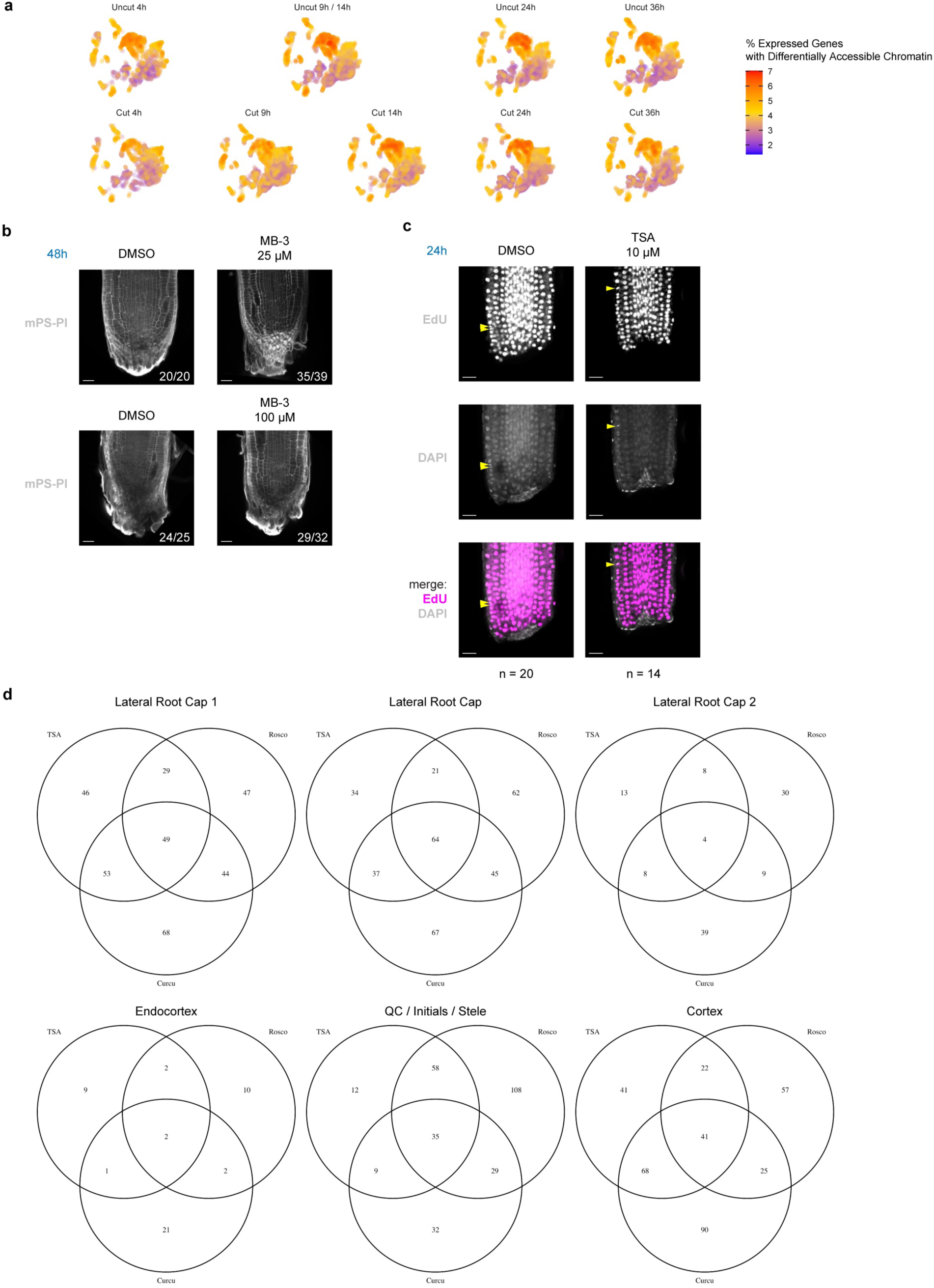
(a) UMAPs showing the relative expression of genes with differentially accessible chromatin in uncut or cut conditions at 4, 9, 14, or 24 hours—uncut 9 and 14 hour timepoints are a shared control. (b) Confocal images of mPS-PI stained roots 48 hours post cut after treatment with mock (DMSO) or 25 µM or 100 µM of the HAT inhibitor MB-3. Fractions indicate the number of observed fully regenerated roots out of total. Scale bars correspond to 20 µm. (c) EdU labeling of mock (DMSO) or TSA-treated roots after 24 hours. Yellow arrowheads indicate mitotic figures. Scale bars correspond to 20 µm. (d) Venn diagrams showing the overlap of genes that are not disturbed by each treatment, i.e. number in the center are the genes never disturbed by any treatment in that cell type. Cell types are the ones where cells in regeneration identified by orange topic are enriched.

**Supplemental Figure 5.**
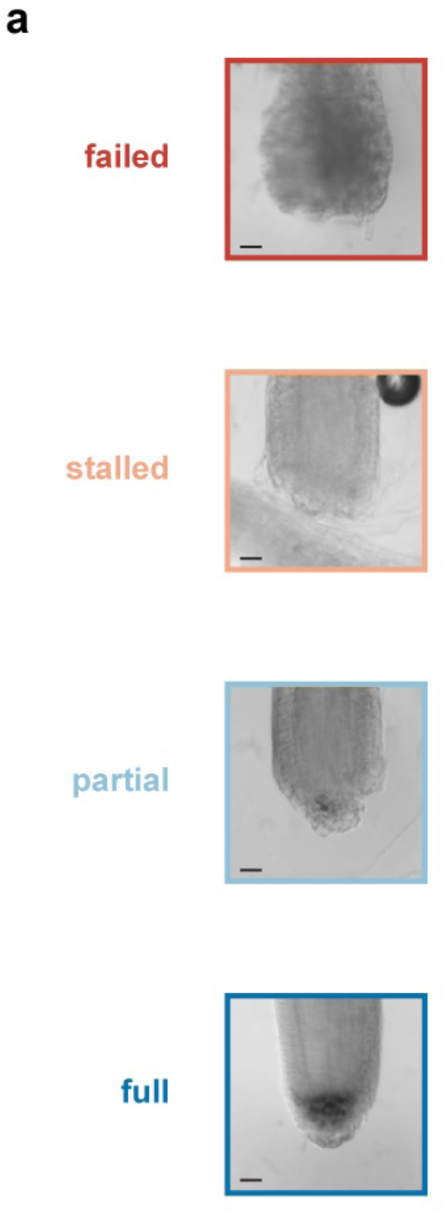
Chromatin accessibility and effects of chemical inhibitors of chromatin modifiers. (a) Phenotypic examples of each regeneration phenotype class seen after pulse-chase experiments treating roots with increasing windows of roscovitine, TSA, and curcumin. Failed roots have expanded root tip cells with emergence of root hairs. Stalled roots are blunt ended and resemble freshly cut roots. Partial roots have some amyloplast-positive cells but lack a fully tapered root tip. Full regeneration entails many amyloplast-positive cells and a continuously tapered root tip. Unscaled images—scale bars estimated from root width and correspond to 20 µm.

**Supplemental Figure 6.**
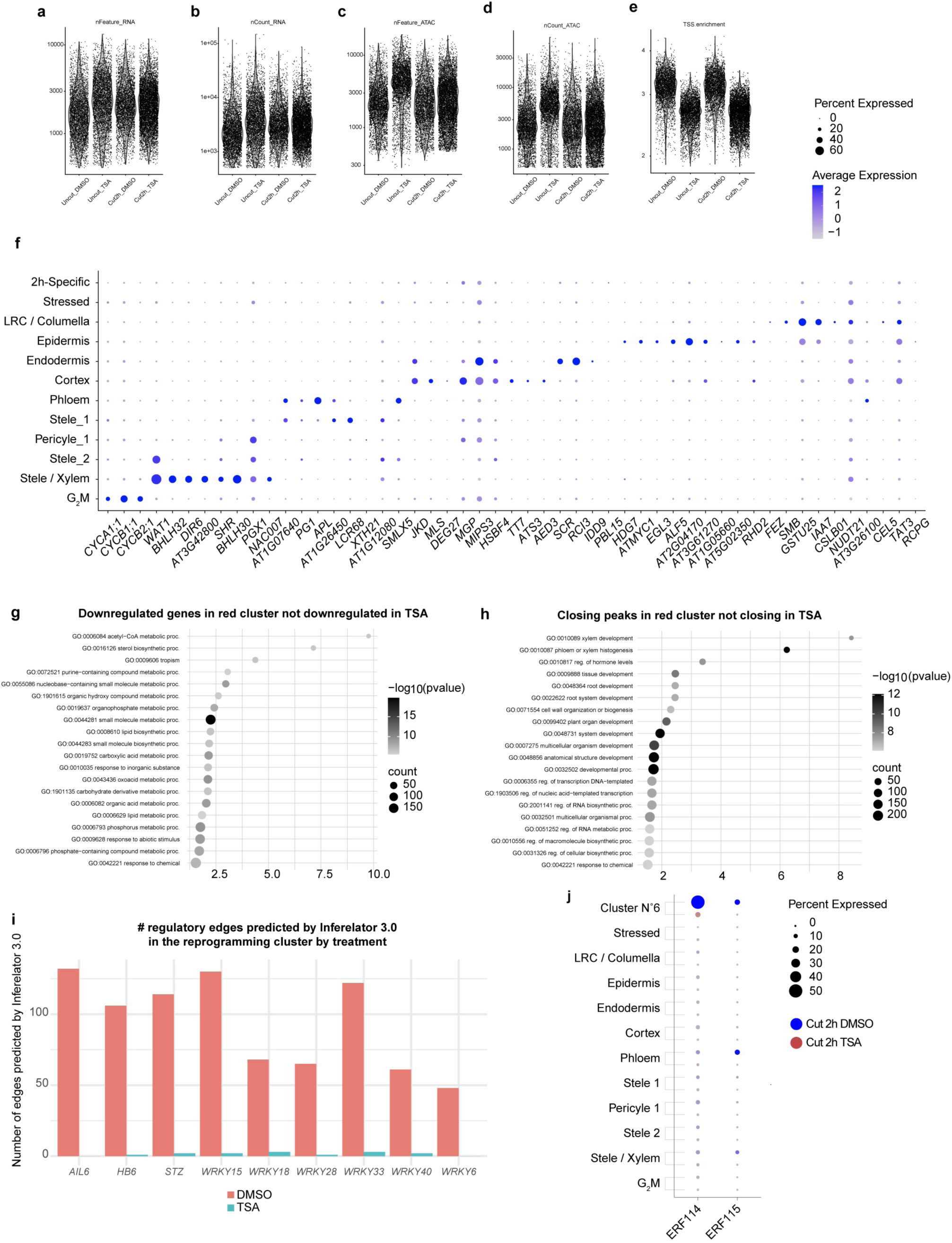
Multiome quality control and gene expression analyses. Multiome graphs showing (a) the number of genes detected per nuclei (b) number of UMI, (c) number of distinct peaks, (d) number of peak UMI, (e) TSS enrichment between the different nuclei and datasets. (f) Dotplot showing cluster-specific gene expression from scRNA-seq data in control cut 0h DMSO condition. (g) Go term enrichment plot of genes that are downregulated in cluster N°6 compared to other cell types in DMSO cut 2h conditions. (h) Go term enrichment plot of genes associated with peaks that are less accessible in cluster N°6 compared to other cell types in DMSO cut 2h conditions. (i) Regulatory edges for regeneration-specific transcription factors were determined with Inferelator 3.0 in the cluster N°6, cut 2h in DMSO and TSA treatment. A confidence cutoff greater than 0.15 was used to indicate a TF-target edge versus non-edge. (j) Dotplot of wound-response genes ERF114 and 115 in control and TSA treated roots, showing expression highly specific to cluster 6 in DMSO and repression in TSA treatment.

**Supplemental Figure 7.**
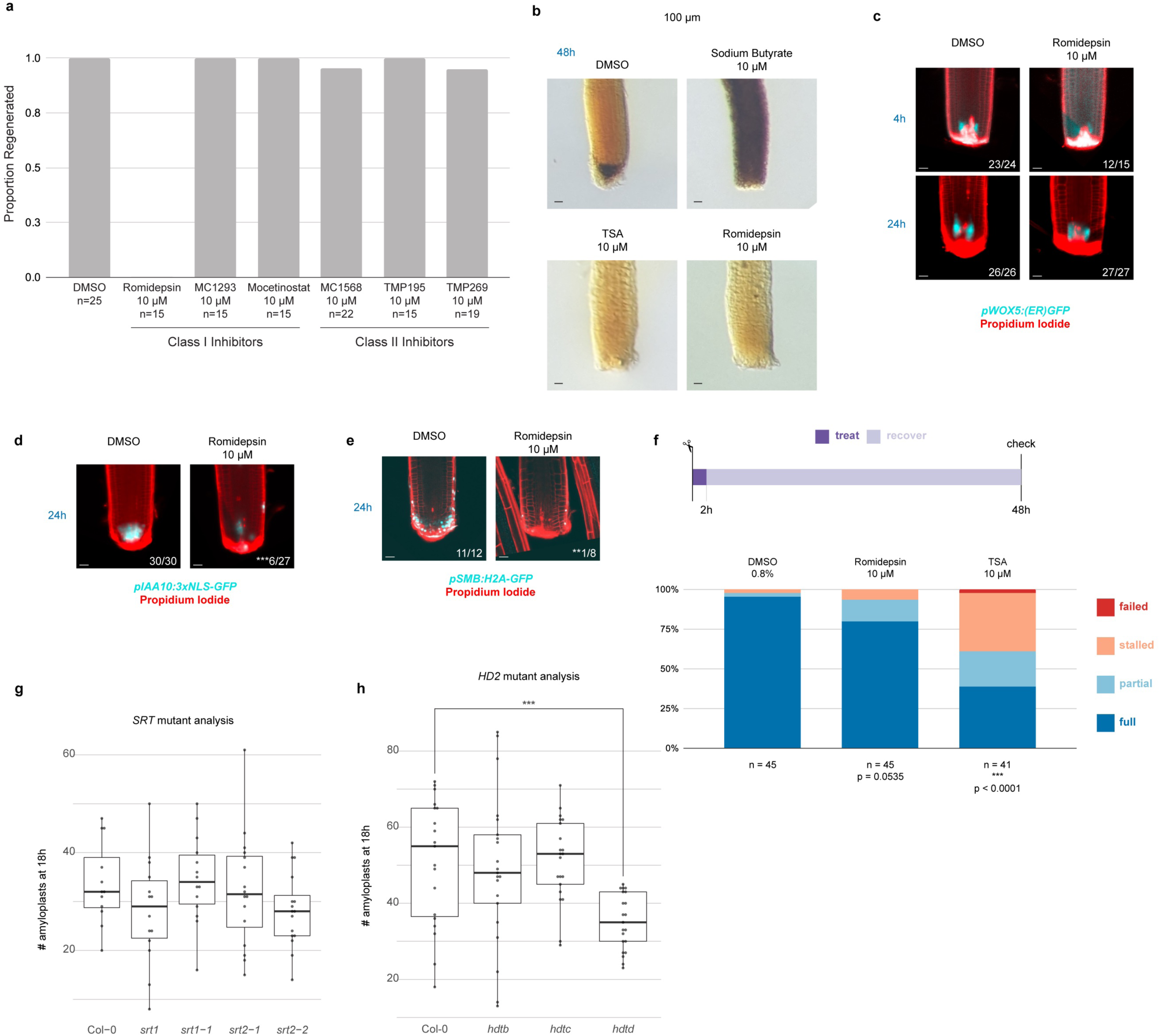
Inhibition of Class I HDACs perturbs regeneration. (a) Results of regeneration assay testing several selective inhibitors of either Class I or Class II HDACs as outlined by Ueda et al. (2017). Only romidepsin, a Class I-specific inhibitor, inhibited regeneration. (b) Lugol staining of amyloplasts in mock (DMSO), sodium butyrate, TSA, or romidepsin treated roots after 48 hours. Unscaled images—scale bars estimated from root width and correspond to 20 µm. (c) Confocal microscopy showing *WOX5* expression after mock (DMSO) or romidepsin treatment at 4 and 24 hours. Fractions indicate frequency of the shown phenotype. Scale bars correspond to 20 µm. (d)-(e) Confocal microscopy showing *IAA10* (d) *or SMB* (e) expression after mock (DMSO) or romidepsin treatment at 24 hours. Fractions indicate frequency of the shown phenotype and asterisks correspond to significance after performing Chi-square with Yates correction. ** = p<0.001, *** = p <0.0001. Scale bars correspond to 20 µm. (f) Top: experimental scheme—plants were cut and treated with mock (DMSO), romidepsin, or TSA for 2 hours before recovery up to 48 hours post cut. Bottom: Stacked bar plot showing fractions of full, partial, stalled, or failed phenotypes for each treatment. P-values derived from Chi-square with Yates correction performed on full regeneration phenotypes vs all other phenotypes between mock and treat. (g) Boxplot showing number of amyloplast-positive cells at 18 hours post cut in wild type (Col-0) or *SRT* mutants *srt1* (SALK_064336C), *srt1-1* (SALK_086287C), *srt2* (SALK_131994C), and *srt2-2* (SALK_149295C) as seen by mPS-PI staining. (h) Boxplot showing number of amyloplast-positive cells at 18 hours post cut in wild type (Col-0) or *HD2* mutants *hd2b* (SALK_113502), *hd2c* (SALK_136925C), and *hd2d* (SALK_095273), as seen by mPS-PI staining. *Hd2d* had significantly fewer amyloplasts compared to wild type, *** corresponds to p=0.0004 from two-tailed t-test after confirming a difference in genotype effect on amyloplasts by ANOVA.

**Supplemental Table 1: scRNAseq analysis of the timepoint**

<u>Sheet1</u>: Summary metrics on scRNAseq datasets

<u>Sheet2:</u>Cell specific marker genes obtained using FindAllMarker function from Seurat V4 using only Control uncut 0h data, raw output.

<u>Sheet3</u>: Pivot table of the different markers, in row genes, in columns cell types and in cells LogFold2.

<u>Sheet4</u>: Curated table of cell type markers genes

<u>Sheet5:</u> Fisher exact test between curated cell type marker genes and upregulated genes between uncut and cut time points for each cell type, this reflects the gain of new identity.

<u>Sheet6:</u> Fisher exact test between curated cell type marker genes and downregulated genes between uncut and cut time points for each cell type, this reflects the loss of identity.

**Supplemental Table 2: Cells in regeneration analysis**

<u>Sheet1</u>:Topic Modelling raw output table per cell

<u>Sheet2</u>: Topic Modelling gene enrichment for each topic

<u>Sheet4</u>: Mini-Ex raw results in the different cell type between CTR (uncut) and CUT conditions

<u>Sheet5</u>: Mini-Ex pivot table analysis per cell type and condition

<u>Sheet6:</u> Raw output data from FindAllMarker function from Seurat V4, differentially expressed genes (DEG) between ctr vs cut, and cell identified as in regeneration.

<u>Sheet7:</u> Curated data frame to identify genes differentially expressed in regeneration.

**Supplemental Table 3: Chemical treatment (TSA, Roscovitin, Curcumin) effect analysis**

<u>Sheet1:</u> Raw output data from FindAllMarker function from Seurat V4, differentially expressed genes (DEG) between ctr vs cut vs all treatments.

<u>Sheet2</u>: Curated DEG table between the different cell type that are enriched for cells in regeneration, in cuts and chemical treatments 14h post cut

<u>Sheet3</u>: Counting table of the number of upregulated and downregulated genes at 14h post cut in DMSO and in the different chemical treatments

<u>Sheet4</u>: Counting table of the number of genes affected by each treatment and their overlaps

**Supplemental Table 4: Multiome Analysis**

<u>Sheet1</u>: Summary metrics on Multiome replicates

<u>Sheet2:</u>Purified marker genes for each cell type in DMSO cut0h condition using theFindAllMarker function.

<u>Sheet3</u>: Purified Peaks for each cell type in DMSO cut0h condition using theFindAllMarker function on the peaks defined by Macs2_peaksNarrowPerCell

<u>Sheet4:</u> Raw output data from FindAllMarker function from Seurat V4, differentially expressed genes (DEG) between cut0h and cut2h and DMSO vs TSA.

<u>Sheet5</u>: Curated DEG dynamic table between the different cell types in cut0h and cut2h and DMSO vs TSA. Values in the table are the avg_log2FC from Tab5.

<u>Sheet6:</u> Raw output data from FindAllMarker function from Seurat V4, Differentially Accessible Chromatin Regions (DACR) between cut0h and cut2h and DMSO vs TSA.

<u>Sheet7</u>: Curated DACR dynamic table between the different cell types in cut0h and cut2h and DMSO vs TSA. Values in the table are the avg_log2FC from Tab7.

<u>Sheet8</u>: Raw output from Motif enrichment function in the different cell type and different treatments.

<u>Sheet9</u>: Curated dynamic table from the Motif enrichment analysis, negative values in the table mean less accessible and positive values mean more accessible.

<u>Sheet10</u>: Mini-Ex raw results in the different cell type between cut0h and cut2h and DMSO vs TSA.

<u>Sheet11</u>: Mini-Ex pivot table analysis per cell type and conditions.

<u>Sheet12</u>: Inferelator output ran on the Cut2h DMSO cluster N°6

<u>Sheet13</u>: Inferelator output ran on the Cut2h TSA cluster N°6

**Supplemental Table 5: GO enrichment and Hypergeometric enrichment of DEG and DACR**

<u>Sheet1</u>: GO enrichment for when comparing uncut vs cut roots, we found that differentially accessible chromatin regions

<u>Sheet2:</u> Hypergeometric enrichment of less accessible chromatin in cluster N°6 and downregulated genes in cluster N°6 with cell specific peaks and cell specific genes.

